# The island rule explains consistent patterns of body size evolution in terrestrial vertebrates

**DOI:** 10.1101/2020.05.25.114835

**Authors:** Ana Benítez-López, Luca Santini, Juan Gallego-Zamorano, Borja Milá, Patrick Walkden, Mark A.J. Huijbregts, Joseph A. Tobias

**Author notes:** These two authors contributed equally (Mark A.J. Huijbregts, Joseph A. Tobias).

## Abstract

Island faunas can be characterized by gigantism in small animals and dwarfism in large animals, but the extent to which this so-called ‘island rule’ provides a general explanation for evolutionary trajectories on islands remains contentious. Here we use a phylogenetic meta-analysis to assess patterns and drivers of body size evolution across a global sample of paired island-mainland populations of terrestrial vertebrates. We show that ‘island rule’ effects are widespread in mammals, birds and reptiles, but less evident in amphibians, which mostly tend towards gigantism. We also found that the magnitude of insular dwarfism and gigantism is mediated by climate as well as island size and isolation, with more pronounced effects in smaller, more remote islands for mammals and reptiles. We conclude that the island rule is pervasive across vertebrates, but that the implications for body size evolution are nuanced and depend on an array of context-dependent ecological pressures and environmental conditions.

## Introduction

From giant pigeons to dwarf elephants, islands have long been known to generate evolutionary oddities^1^. Understanding the processes by which island lineages evolve remains a prominent theme in evolutionary biology, not least because they include many of the world’s most bizarre and highly threatened organisms^2^. The classic insular pattern of both small-animal gigantism and large-animal dwarfism in relation to mainland relatives has been described as a macro-evolutionary or biogeographical rule – the ‘island rule’^3–5^ (Fig. 1). However, previous research has cast doubt on the generality of this pattern^6^, suggesting that body size shifts are asymmetrical, with reduced size in some clades (e.g. carnivores, heteromyid rodents, and artiodactyls) or increased size in others (e.g. murid rodents)^7, 8^. Even in these cases, the underlying mechanisms driving patterns of insular gigantism and dwarfism remain unclear.

**Figure 1.**
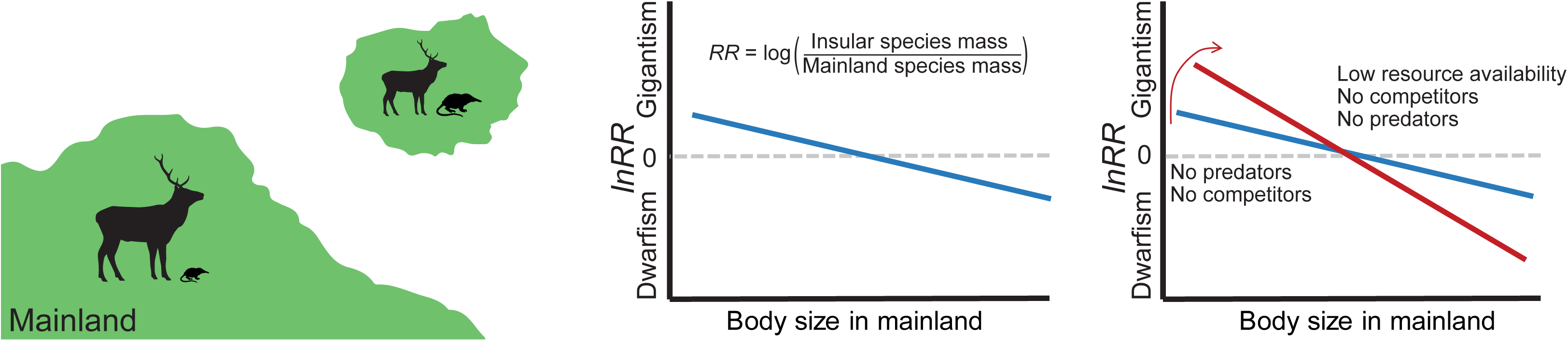
Conceptual figure showing body size evolution in island populations. According to the island rule, changes in body size of island populations are dependent on the body mass of mainland relatives, with small species tending to increase in size on islands (gigantism) and large species tending to decrease in size (dwarfism). By plotting the log response ratio (*lnRR*) between insular mass and mainland mass, against mainland mass, we can test if insular populations adhere to the rule (intercept > 0 and slope < 0) (blue line). Mechanisms proposed to drive ‘island rule’ effects are mainly based on reduced predation, inter- and intra-specific competition, and food availability, suggesting that the relationship will steepen in small, remote islands (red line).

Several mechanisms have been proposed to explain the island rule, including reduced predation, relaxed competition and resource limitation in island environments^9^. In theory, each of these factors may be accentuated in smaller, more isolated islands, where lower levels of interspecific competition and predation could lead to ‘ecological release’, allowing small-bodied species to increase in body size^5, 9^. Conversely, among large-bodied species, limited resource availability could select for smaller body sizes with reduced energy requirements, leading to insular dwarfism. Climatic conditions may also influence body size evolution on islands since primary productivity and associated resource availability are strongly influenced by climate^9, 10^. Although previous studies of body size evolution on islands have tested the effects of these different mechanisms, many have focused on relatively restricted geographic and taxonomic scales and did not directly address the island rule in its broad sense across multiple species within a taxon^10–13^, with notable exceptions^9, 14–16^.

Most work on the island rule has been restricted to mammals (e.g.^4, 7, 14, 17^), although the hypothesis has also been tested in amphibians^18^, reptiles^19–21^, birds^15, 22^, fish^23^, insects^24^, molluscs^25^, and plants^26^. The highly inconsistent results of these studies (e.g.^5, 6, 27^) are perhaps unsurprising because they typically deal with single species or pool together data on different traits from numerous sources without controlling for variation in study design or accounting for sampling variance. Accordingly, a recent systematic review based on a simplified scoring system^27^ concluded that empirical support for the island rule is not only potentially biased but also generally low, particularly for non-mammalian taxa. However, scoring approaches provide only limited information as they do not account for heterogeneity between studies, taxonomic representativeness, sample size, or precision in the estimates.

These limitations are best addressed with formal meta-analyses^28, 29^, hence we tested the island rule hypothesis by applying phylogenetic meta-regressions to a global dataset of 2,479 island-mainland comparisons for 1,166 insular and 886 mainland species of terrestrial vertebrates (Supplementary Dataset 1, Fig. 2). Our analytical framework allows us to control for multiple types of variation, including data source, sample size imbalance, intraspecific and intra-population variability, and phylogenetic relatedness (see Methods). For each island-mainland comparison, we calculated the log response ratio (*lnRR*) as the natural logarithm of the ratio between the mean body size of individuals from an insular population *M_i_* and that of mainland relatives *M_m_* (*lnRR* = log[*M_i_/Mm*])^30^. Then, we regressed *lnRR* against the body mass of the mainland population (*M_m_*)(Fig.1).

**Figure 2.**
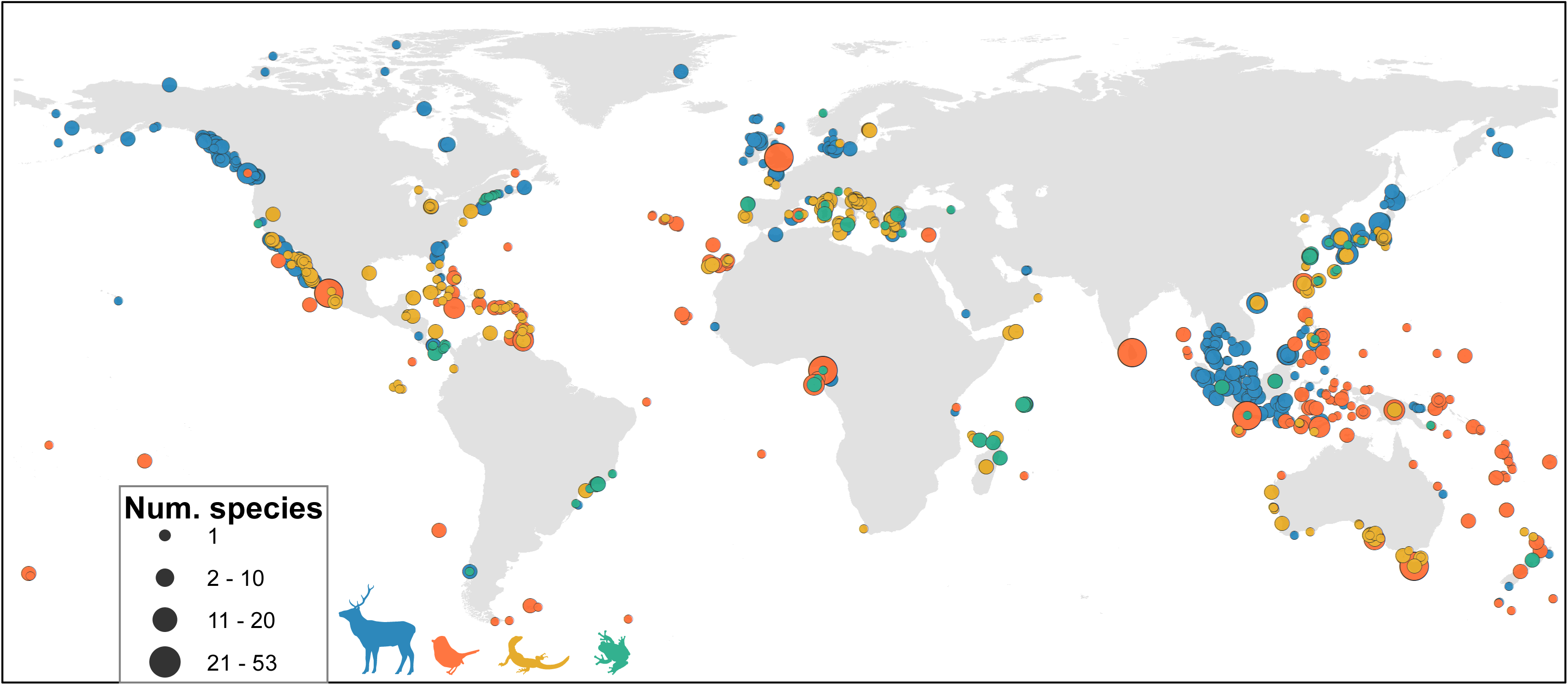
Location of island populations included in our analyses for mammals (N = 1058, blue), birds (N = 695, orange), reptiles (N = 547, yellow), and amphibians (N = 179, green). The size of each point indicates the number of species sampled on each island; some points overlap. See Fig. S1 for a 4-panel figure with the location of insular populations separated for each taxonomic group.

This framework provides a clear set of predictions in the context of evolutionary trajectories on islands^4, 6, 14^. Specifically, since negative values of *lnRR* indicate dwarfism and positive values indicate gigantism, a positive intercept and negative slope of the *lnRR-*mainland mass relationship supports the island rule (Fig. 1). Given the contentiousness of the generality in the island rule, we assessed the robustness of our results against potential biases derived of regressing ratios^31, 32^, using small samples, imputing missing data, or the influence of using data from the island rule literature or derived from other studies focused on unrelated questions (i.e. publication bias; see Methods). Finally, we use our framework to assess how body size shifts are related to island size, island isolation, island productivity and climate, as well as species diet. The extent to which these different factors explain insular body size shifts allows us to re-evaluate a range of hypotheses for the mechanisms underlying “island rule” effects on body size, including ecological release^9^, immigrant selection^9^, resource limitation^9, 33, 34^, thermoregulation^9, 15, 35^, water availability^36, 37^ and starvation resistance^9, 33^ (Supplementary Table 1, Extended Data Fig. 1).

## Results

### The generality of the island rule

We found that *lnRR* and mainland body mass were negatively related for mammals, birds and reptiles, with small species tending to gigantism and large species to dwarfism (Fig. 3). The relationship was weakly negative but statistically non-significant for amphibians, with a tendency towards gigantism across all body sizes (Fig. 3, Table 1). We obtained similar results using size ratios corrected for small sample size (*lnRR*^Δ^), or by regressing island mass against mainland mass, with support for the island rule across all groups except for amphibians (Supplementary Table 3-4). This indicates that our analyses are robust to small sample size bias^38^ or any potential spurious correlation associated to ratio regression models^31, 32^ (Extended Data Fig. 2). Further, neither imputation nor publication bias influenced our results (Supplementary Table 5-6), with no apparent differences between island-mainland comparisons sampled from studies formally testing the island rule or compiled from unrelated data sets.

**Figure 3.**
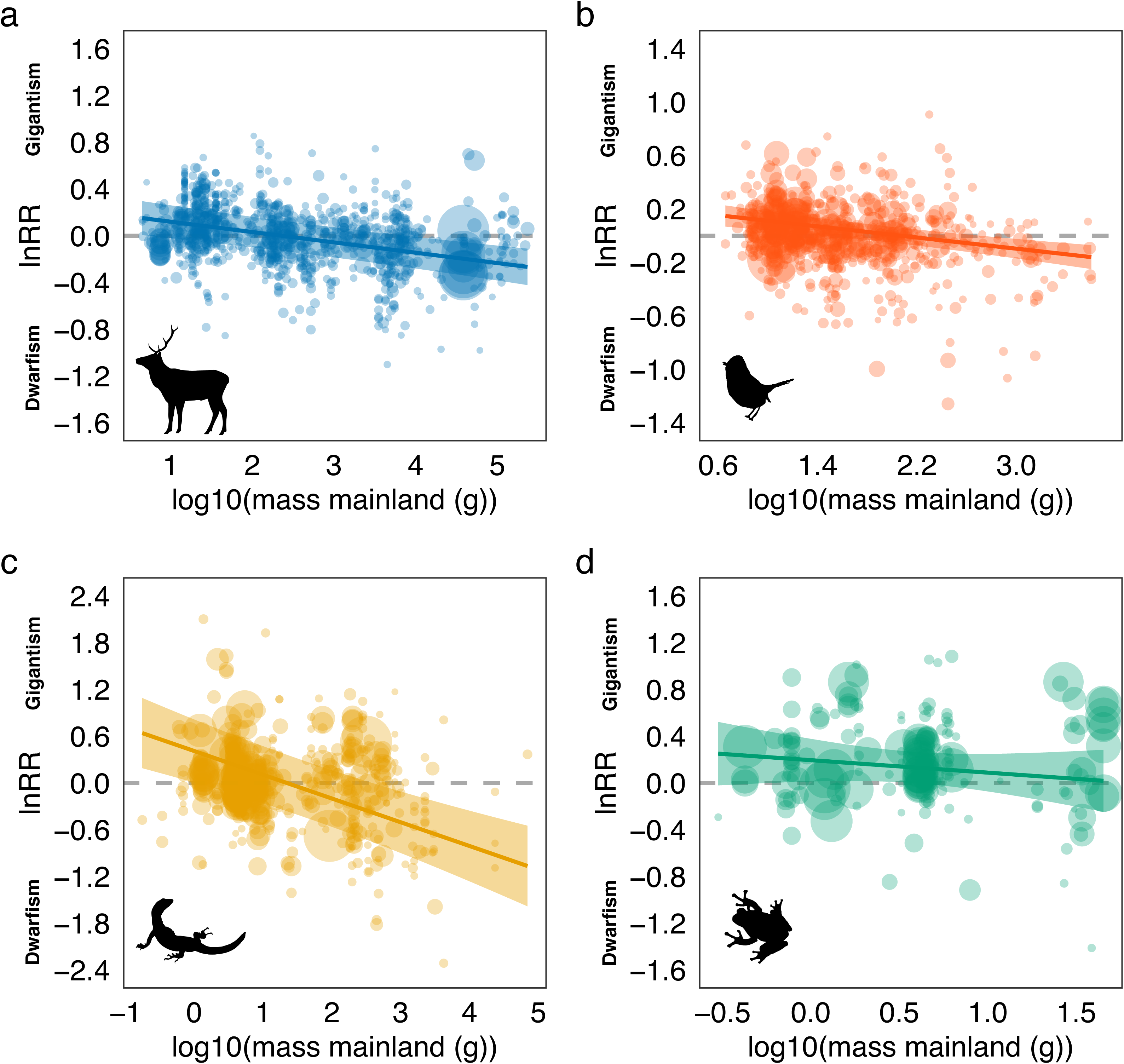
Relationship between *lnRR* (log response ratio between island mass and mainland body mass) and body mass in the mainland for (a) mammals (N = 1058), (b) birds (N = 695), (c) reptiles (N = 547) and (d) amphibians (N = 179). Models were fitted using phylogenetic multi-level meta-regression models with mainland body mass as moderator, and observation-level ID, source ID, species ID and phylogeny as random effects. *lnRR* > 0 indicates gigantism; *lnRR* < 0 indicates dwarfism; and *lnRR* = 0 indicates stasis (no shift in body size from mainland to island populations). The size of points represents the inverse of the sampling variance for each paired island-mainland response ratio in the model. Shaded areas represent 95% confidence intervals. Note that y-axes have different scales.

**Table 1.** Parameter estimates for the phylogenetic meta-regression models testing the generality of the island rule in terrestrial vertebrates. *k*: number of island-mainland comparisons (*lnRR*), *Q_m_*: test of moderators (log_10_(mainland mass). *R^2^*m* :* marginal *R^2^*, estimated percentage of heterogeneity explained by the moderator (fixed effects). *R^2^*c* :* conditional *R*^2^, percentage of heterogeneity attributable to fixed and random effects.

| Class | $k$ | Intercept<br>(CI) | Slope<br>(CI) | $Q_m$<br>(p-value) | $R^2_m$ | $R^2_c$ |
| --- | --- | --- | --- | --- | --- | --- |
| Mammals | 1058 | 0.208<br>(0.052 – 0.365) | -0.088<br>(-0.122 – -0.055) | 27.30<br>($p < 0.001$) | 11.4 | 56.4 |
| Birds | 695 | 0.216<br>(0.117 – 0.315) | -0.104<br>(-0.145 – -0.064) | 25.40<br>($p < 0.001$) | 7.0 | 43.4 |
| Reptiles | 547 | 0.410<br>(0.006 – 0.814) | -0.305<br>(-0.419 – -0.190) | 27.21<br>($p < 0.001$) | 17.6 | 66.6 |
| Amphibians | 179 | 0.195<br>(0.012 – 0.377) | -0.107<br>(-0.320 – 0.107) | 0.96<br>($p = 0.328$) | 1.4 | 67.1 |

Mainland body mass explained 11.4, 7.0 and 17.6% of the variance in mammals, birds and reptiles, respectively. The amount of further variance accounted for by phylogeny (0.0–29.8%), data source (1.8–25.1%), and species (25.9–53.2%) fluctuated widely among taxa (Extended Data Fig. 3). Phylogeny accounted for a relatively large amount of variance in mammals (20.1%) and reptiles (29.8%), but even in these cases the overall patterns were not driven by large effects in particular clades. Some groups tended towards gigantism and others towards dwarfism, while others contained both dwarfs or giants depending on body size (e.g., Primata, Rodentia, and Carnivora in mammals, and Viperidae, Scincidae and Iguanidae in reptiles; Extended Data Fig. 4).

**Figure 4.**
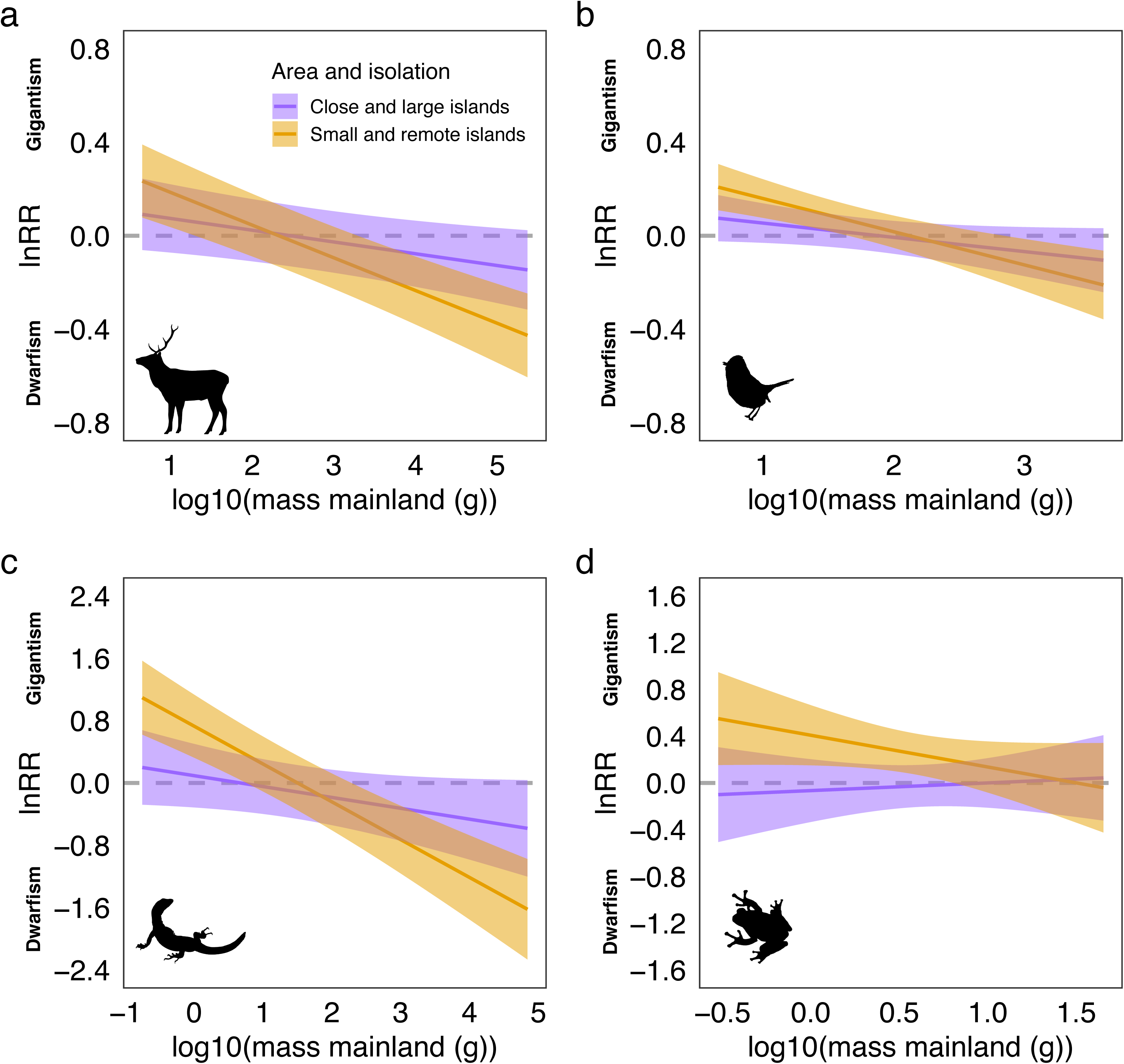
The effect of island area and spatial isolation on insular size shifts in terrestrial vertebrates for (a) mammals (N = 1058), (b) birds (N = 695), (c) reptiles (N = 547) and (d) amphibians (N = 179). Continuous variables are represented at the 10% and 90% quantile for each extreme (close vs remote islands; small vs large islands). *lnRR* > 0 indicates gigantism; *lnRR* < 0 indicates dwarfism; and *lnRR* = 0 indicates stasis (no shift in body size from mainland to island populations). Shaded areas represent 95% confidence intervals.

### Ecological mechanisms underlying body size evolution on islands

The pattern of body size evolution in our island-mainland comparisons provides some insight into the likely mechanisms driving “island rule” effects (Extended Data Fig. 5-8, Supplementary Table 7, Supplementary Dataset 2). Overall, insular size shifts arise through some combination of ecological release from predation and competition, resource limitation, biased colonization (i.e. immigrant selection), and starvation resistance. The fact that no single factor explained island effects on body size is not surprising because some hypotheses shared overlapping predictions, making them difficult to disentangle.

Shifts in body mass of mammals were mostly explained by island size and spatial isolation (*Q_m_* = 12.20, P = 0.002, Fig. 4a), resulting in more pronounced gigantism or dwarfism in small and remote islands. Birds showed similar size shifts in relation to spatial isolation and island area, but these were not statistically significant (Supplementary Table 7). In both mammals and birds, temperature had similar effects across the size range, with body size consistently larger in cool islands and smaller in warm islands (Extended Data Figs. 5e, 6e). Hence, in these groups, even large species that had undergone dwarfism were significantly larger in cool insular environments than in warm ones. Contrary to the starvation resistance hypothesis, small-sized birds did not become larger in highly seasonal islands, but large-sized birds had reduced dwarfism on islands with high seasonality in temperatures (*Q_m_* = 12.33, P < 0.001, Extended Data Fig. 6).

In reptiles, the combination of island area and spatial isolation were the most important factors explaining variation in body size (Fig. 4c), with productivity and seasonality also supported but with weaker effects (Extended Data Fig. 7). Similar to mammals, the tendency towards dwarfism or gigantism in large-bodied or small-bodied reptiles was more apparent in isolated small-sized islands, with stronger effects of area than isolation (Supplementary Table 7). The effects of productivity and seasonality were only partially in line with predictions, as small-sized species were larger on islands with high seasonality, but smaller on islands with high productivity (Extended Data Fig. 7). In turn, large-bodied reptiles were smaller on islands with low productivity and high seasonality.

Finally, the relationship between size ratio and mainland mass in amphibians was slightly steeper in small and remote islands (Fig. 4d), with island area being marginally more important than spatial isolation (Extended Data Fig. 8). The effect of seasonality was clearer, with amphibian species inhabiting islands with high seasonality (unpredictable environments) tending toward gigantism, whereas those from islands with low seasonality (predictable environments) being similar in size to mainland counterparts (Extended Data Fig. 8). We found no effects of diet for any of the four taxa, or precipitation for amphibians, contrary to the water availability hypothesis.

## Discussion

Based on comprehensive morphometric data from a worldwide sample of island fauna, we show consistent patterns of body size evolution across terrestrial vertebrates in accordance with predictions of the island rule. This finding was robust to alternative modelling approaches (island mass vs mainland mass regressions), small sample bias, data imputation, and publication bias. Moreover, we have demonstrated that insular size shifts are contextual and depend not only on the body size of mainland relatives (island rule *sensu stricto*) but also on the physiographic and climatic characteristics of particular island environments^9^.

### Repeated evolutionary trajectories on islands

We found a clear negative relationship between insular body size variation and the body mass of mainland individuals in mammals, birds, and reptiles. Mainland body mass explains between 7.0 and 17.6% of the variation in insular size divergence in these three taxonomic groups, which is similar to that reported in smaller-scale studies of bats (15%), birds (13%), snakes (42%), non-volant and terrestrial mammals (11–21%), and turtles (8%)^5, 14, 15, 39, 40^. Contrary to these earlier studies, our analyses are corrected not only for phylogenetic relatedness, but also for variability between species and intrapopulation variability, thereby strengthening the evidence for predictable evolutionary trajectories on islands. Nevertheless, the island rule provides only a partial explanation for these trajectories because substantial variation around the trend line remains unexplained. We also conducted the first multispecies test of island rule effects in amphibians, showing that the relationship goes in the expected direction but with a weak effect (1.4 %), possibly because the body mass range in amphibians is narrower and limited to small sizes (∼ 0.5-50 g) and thus most amphibians tend to gigantism on islands with reduced predation risk.

Our findings are in contrast with a number of studies rejecting the island rule, including a recent review of evidence from across mammals, birds and reptiles^27^, as well as other taxon-specific studies focused on lizards^20, 41^ and turtles^21^. On the other hand, the patterns we detect are consistent with analyses supporting the island rule in snakes^19^, mammals^4, 9^ and birds^5, 15^ We conclude that the contradictory results of previous studies may have been related to sampling bias, heterogeneity between sources and species, and phylogenetic relatedness (i.e. statistical non-independence). By accounting for these effects in our global models we are able to demonstrate that vertebrate animals evolve in largely consistent ways on islands. Further, we have shown that the island rule is not clade-specific and instead applies to numerous clades within major taxonomic groups, particularly in mammals and birds.

A corollary that emerges from the island rule is that body size converges on islands. Specifically, if insular environments select for intermediate body sizes, closer to the optimal size of the focal clade, then the size spectrum of organisms found on islands should be narrower compared to the mainland^42, 43^. Theoretically, the optimal body size towards which small and large species may converge in low-diversity systems such as islands should correspond to the point where the trend intersects the horizontal dashed line in the relationship between size ratio and mainland mass, at which point fitness is maximized^42^ (but see^44^). Interestingly, the shift between dwarfism and gigantism in our models occurred at approximately 100-250 g in endotherms, slightly larger than the 100 g adult body mass proposed for mammals^42^ (but see^43^), and the mode of the global body size distribution of birds that separate between small- and large-bodied species (60 g)^22, 45, 46^. Additionally, our analyses suggest that the optimal body size for island reptiles should be ca. 20 g, which is marginally higher than the modal body size of Lepidosaurs (14.1 g)^47^. Whether there is an optimal body size in island biotas has been the subject of much debate^44^, but overall we expect that phenotypic variability in morphometric traits will be substantially narrowed if directional selection is operating in island assemblages, a feature that warrants further investigation. Additionally, optimal phenotypes should vary with the environmental characteristics of islands, in particular their area and isolation, climate, productivity and seasonality. For example, in mammals, our results suggest that the optimal body size would be ca. 100 g and ca. 900 g in warm and cold islands, respectively.

### Ecological mechanisms influencing body size variation

Because body size is intimately linked to many physiological and ecological characteristics of vertebrates, it may be associated with a variety of environmental factors. As a consequence, the body size of colonizing species may predictably evolve as the result of selective pressures associated with insular environments (e.g., low food resources, few competitors, no predators) and others that act across larger geographic scales (e.g., climate). For mammals and reptiles, our results suggest that insular body size shifts are indeed governed by spatial isolation and island size, with individuals becoming dwarfs or giants in remote islands of limited size. Furthermore, the slope of the relationship between size ratio and mainland mass was slightly steeper for birds and amphibians in small remote islands than in large islands near continental land masses (Fig. 4). This points to a combination of resource limitation (with small islands having fewer resources to maintain large-sized organisms^48, 49^) along with release from interspecific competition and predation pressure in small, species-poor islands. The pattern is also consistent with biased colonization favoring larger individuals with higher dispersal abilities (immigration selection^50^). Conversely, our results showed that body size divergence on islands close to the mainland was minimal, reflecting two non-mutually exclusive processes. First, many of these islands were connected to the continent by land bridges so recently that phenotypic differences have not had time to accumulate. Second, regular dispersal between mainland and island populations promotes gene flow, with introgression counteracting divergent selection^51, 52^.

Besides island physiography (area and isolation), other relevant factors were temperature conditions in endotherms and resource availability and seasonality in ectothermic organisms. Mammals and birds both responded to island temperature in line with the heat conservation hypothesis, with small- and large-sized species exhibiting exacerbated gigantism and diminished dwarfism, presumably to conserve heat in colder, harsher insular environments. Additionally, temperature seasonality was an important determinant of the size of large-bodied birds, with populations on highly seasonal islands being similar in size to mainland populations. One possibility is that larger size in these cases may help maintain energy reserves during periods of low food availability, allowing them to thrive in otherwise hostile environments. Another possibility is that bird populations on highly seasonal islands – which tend to be situated at relatively high latitudes – are more often seasonally mobile or even migratory, potentially increasing gene flow with mainland populations or weakening adaptation to the local environment^53^. These findings add new insights to previous results regarding the role of thermal and feeding ecology on morphological divergence in island birds^54, 55^. Traditionally, changes in feeding ecology were thought to be the prime force in driving morphological divergence in island birds^54, 55^. Yet, our results imply that physiological mechanisms related to heat conservation (‘thermoregulation hypothesis’) and energy constraints (‘starvation resistance hypothesis’) may also shape body size evolution in island birds.

In reptiles, we find some evidence that resource availability and seasonality are important factors explaining body size evolution, with some deviations from the patterns predicted. As hypothesized, large species are much smaller on islands with low resource availability, and small species are larger on islands with high seasonality. Yet, unexpectedly, small species are larger on islands with low productivity, perhaps because increased intraspecific competition favors large individuals under the high population densities that reptiles often attain on islands^56, 57^.

Overall, most amphibians tended to gigantism, presumably as a result of increased growth rate or lower mortality due to reduced predation pressure on islands^58^. Additionally, we found that body size of amphibians consistently increased on islands where resources were highly seasonal and unpredictable, perhaps to maximize energy reserves and withstand long periods without food, for example during aestivation or hibernation^59^. We did not find a clear relationship between precipitation and body size, suggesting that water availability is not a key factor. It appears that gigantism in island amphibians is mostly driven by physiological mechanisms that maximize growth rate, particularly in smaller, more isolated islands. These findings should be further explored when more data on island-mainland pairwise populations of amphibians become available.

### Body size evolution in extinct species

Our analyses focused on extant species for which we could gather information on the variation around the morphometric estimates, along with sample size (essential for meta-analyses). The widespread extinction of large species on islands, including dwarf morphotypes of large species such as insular elephants in Sicily and the Aegean islands^60, 61^, may have masked the historical pattern of phenotypic variation on islands^62^. Giant insular birds^54, 63^, primates^64, 65^, and lizards^66^, along with large insular turtle species, went extinct during the Holocene and late Pleistocene^67^, most likely because of overhunting and the introduction of invasive species^68, 69^. Overall, it is estimated that human colonization of oceanic islands was followed by the extinction of 27% of insular endemic mammals^70^, as well as over 2000 bird species in the Pacific region alone^71^, with these losses biased towards large-bodied, flightless, ground nesting species^68^. Extinct species may shed new light on size evolution in insular vertebrates because species extinctions have substantially altered the biogeography of body size in island faunas, potentially leading to downsized insular communities^72, 73^. For example, the predominance in our dataset of smaller-bodied organisms could reflect the extinction of large species on islands^68^, or simply the fact that few islands support large species. Either way, further studies should include data from extinct species as this may alter or strengthen the signal that we report for extant species^39^.

We foresee that, under global change, the extinction of insular species and the introduction of novel (invasive) species may trigger new equilibria, with concomitant shifts in the composition of insular communities and the opening of novel niches to which species may respond via genetic adaptations and phenotypic plasticity. Recent evidence indicates that even introduced species on islands, which were not included in our analysis, predictably evolve towards dwarfism or gigantism^74–76^. In theory, as the Anthropocene gathers pace, further extinctions will drive a decline in mean body size of the overall island community, pushing optimal body sizes towards the lower end of body size ranges in the different vertebrate groups.

### Conclusions

Of the many evolutionary implications of living on islands – together known as the ‘island syndrome’^2^ – the effects on body size are the most widely known and controversial. We have shown that these ‘island rule’ effects are widespread in vertebrate animals, although the evidence for amphibians is inconclusive. Morphological changes were directional for species at the extremes of the body size range in mammals, birds and reptiles, following the predicted pattern of convergence towards intermediate “optimum” body sizes, in line with optimal body size theory^42, 43, 45^. Although this convergence towards morphological optima may result from natural selection or phenotypic plasticity, the exact mechanism producing these changes on islands is still not well understood. Nonetheless, we found that consistent transitions towards intermediate body sizes were associated with a combination of factors, indicating a range of different ecological mechanisms. Our results highlight the contextual nature of insular size shifts, where island physiographic, climatic and ecological characteristics play a fundamental role in shaping body size evolution, reinforcing the idea that large-scale macroevolutionary patterns do not arise from single mechanisms but are often the result of multiple processes acting together^77, 78^.

## Methods

### Data collection

We collected baseline morphometric data from articles included in a recent assessment of the island rule^27^, as well as other compilations assembled to test the hypothesis in reptiles^20^, mammals^6^, and birds^15^. To expand this sample, we then performed a literature search (February 2020) in Web Of Science Core Collection (WOS) using the following search string: (“island rule” OR “island effect” OR “island syndrome” OR island*) AND (gigantism OR dwarfism OR “body size” OR weight OR SVL OR snout-vent length OR length OR size) AND (mammal* OR bird* OR avian OR amphibia* OR reptile*) (Appendix 1). Because this search was complementary to the data we have gathered from previous compilations^6, 15, 20, 27^, we only downloaded the first 500 hits out of a total of 33,431 hits ordered by relevance, and removed duplicates already included in our dataset. We reviewed every island-mainland comparison reported in published studies and traced primary source data when possible to extract original measurements. We also extracted data from all studies containing morphometric measurements for insular populations when these could be matched with equivalent data published elsewhere for relevant mainland taxa. We excluded problematic data, such as comparisons that were not supported by taxonomic or phylogenetic evidence, or which reported morphometric data restricted to single specimens or without sample size. In addition, we excluded comparisons based on extinct taxa since they are often known from very few or incomplete specimens (Supplementary Dataset 3).

It has been argued that research on the island rule might be prone to ascertainment bias, where researchers are more likely to notice and measure animals of extreme body size when conducting research on islands^41^. To help overcome this problem, we collected body size data not only from studies testing the island rule, or reporting dwarfism and gigantism in island fauna, but also from studies that did not specifically test hypotheses related to the island rule. We matched unpaired insular populations with independent data from mainland populations by performing species-specific searches in WOS and Google Scholar. We also compiled morphometric data for 442 insular and 407 mainland bird species from an independent global dataset of avian functional traits^79^.

Large islands may be more ‘mainland like’ in relation to factors that are thought to affect body size (i.e. competition, resource availability and predation^5^). Thus, when major islands were at least 10 times larger than a nearby island, we treated the large island as the mainland comparison, following previous studies testing the island rule^4, 5, 20^. Consequently, a single mid-sized island can simultaneously be treated as the continent in comparisons with smaller islands, and the island in comparisons with larger continents. When authors reported data referring to an entire archipelago instead of a specific island (3.2% of cases), we used the size of the largest island as island area. Removing these cases from our analyses did not qualitatively affect our results (Supplementary Table 8).

Our final dataset contained 529 data sources and 2,479 island-mainland comparisons^7, 10, 36, 58, 79–602^. In total, we collated morphometric measurements for 63,561 insular and 154,875 mainland specimens representing mammals (1,058 island-mainland comparisons), birds (695 comparisons), reptiles (547 comparisons) and amphibians (179 comparisons) from across the globe (Fig. 2). 2,068 island-mainland comparisons (83.4%) were within species (e.g. subspecies) comparisons, and 411 (16.6%) were between-species comparisons. Insular populations were sampled from an array of islands varying widely in size (0.0009–785,753 km^2^), climate, and level of spatial isolation (0.03–3,835 km from mainland). To explore the drivers of body-size shifts in insular populations, we also sampled species with a wide range of average body masses (0.18–234,335 g). We collated data on body size indices (body mass, body length, cranial and dental measurements) of different taxa in island and mainland populations following strict morphological, phylogenetic and biogeographic criteria. Specifically, we always compared the same body size index for island and mainland populations. For within-species comparisons, we compared island and mainland populations based on the information given by the authors of the relevant study (e.g. taking note of which mainland source populations are likely to inhabit a particular island because of colonization history or isolation via rising sea levels^89, 101, 240, 386, 548^). When we matched comparisons independently, we used information published in the study reporting the insular form, selecting the geographically closest mainland population whenever possible. In addition, we prioritized latitudinal alignment of mainland and island populations to avoid confounding effects of latitudinal variation in body size. In the case of island endemics, we compared island populations to their closest mainland relative whenever these were identifiable by phylogenetic data or other information reported in each particular study. This usually meant selecting their sister species or the geographically closest representative of a sister clade or polytomy (Supplementary Dataset 1). If we could not reliably establish the closest mainland relative, we discarded the data (see Supplementary Dataset 3).

When more than one body size index was reported in published studies, we prioritized those indices related most closely to body mass (Supplementary Table 2). For mammals, we selected indices in this order of preference: body mass, body length, cranial length (greatest skull length or condylobasal length), and dentition (e.g. canine length)^5^. For birds, preferred indices were body mass, wing length, tarsus length and bill length. Finally, for amphibians and reptiles, size was reported as body mass, snout-vent-length (SVL), carapace length (CL, for turtles) and total length (TL, including SVL and tail length). In all cases, we included measurements for adults only. To avoid size biases attributable to sexual size dimorphism, we calculated the pooled mean for both sexes and the combined SD using standard formulae for combining groups^603^. When information was only available for one sex (male or female), we restricted our size comparisons to the sex for which we had morphometric data in both mainland and island populations. Data from zoos or studies that could not be georeferenced were discarded.

To overcome the problem that different authors report size using different indices, we used allometric relationships to convert island and mainland size to body mass equivalents, thereby enabling cross-taxa and cross-study comparisons. Although this conversion is imprecise, morphological indices and body mass are nonetheless highly correlated across the global scale and wide range of body sizes within our samples (providing more accurate predictions than simply assuming an exponent ∼3, as in previous studies testing the island rule^5, 9^). We used published allometric relationships where available (see Supplementary Table 2), or derived them based on published datasets^47, 179, 604–609^ and other data sources (Supplementary Dataset 4). To calculate allometric relationships, we used OLS (Ordinary Least Square) models of the log^10^ transformed body mass against the log^10^ transformed body size index (e.g. condylobasal length, Supplementary Table 2, Supplementary Dataset 4).

For birds, we complemented published data with standardized morphometric measurements from 3,618 museum specimens and live individuals of 436 insular and 404 mainland bird species (see^79^). We used wing length in the main analyses instead of tarsus length because the former is a better predictor of body mass in our dataset (*R^2^_wing_* = 0.89 vs *R^2^_tarsus_* = 0.69, Supplementary Table 2) (see also^610^). Although wing length may change during moult or thereafter because of wear, these effects are negligible in relation to interspecific differences^79^ and minimized by calculating averages across multiple individuals. Further, interobserver differences between measurements may explain some variation in wing length estimates, but again this bias was shown to have negligible effects in our dataset by comparing repeated measures from different observers (see^79^). To assess the consistency in our results, we repeated analyses using tarsus length, another popular proxy of overall body size in birds^611^. Our results were unchanged (Fig. S2).

To select suitable comparisons for museum specimens, we first classified species as either insular or continental by overlapping IUCN range polygons with a GIS land layer including continental land masses. For each insular species we then identified continental sister species from avian phylogenies^612^, using the method described above. We excluded bird species that are highly pelagic or aerial (e.g. swifts) and fully migratory species because in these groups it is unclear whether insular and mainland forms experience different environments^15^. Further, we also excluded flightless bird species, because morphological changes may be due to flightlessness rather than island dwelling per se^15^.

We calculated the response ratio (*lnRR,* eq. 1) as effect size in our meta-regressions, where we divided the mean body mass of individuals from an insular population 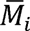 by that of the nearest mainland relative, 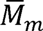, and then applied the natural logarithm. Unlike unlogged ratios, the sampling distribution of *lnRR* is normal, particularly for small samples^30^, and thus less prone to statistical artefacts associated with ratio-based regressions.

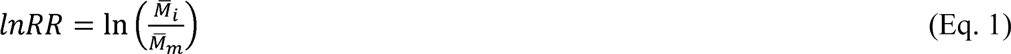

Response ratios greater than zero indicate a shift towards larger sizes (gigantism) whereas ratios less than zero indicate shifts towards smaller sizes (dwarfism). Besides mean measurements, we recorded measures of variation, i.e. standard deviation (SD), standard error (SE) or coefficient of variation (CV), and sample sizes of the body size indices in island and mainland organisms. SD and sample sizes were used to calculate sampling variances (Eq. 2), which were then used to weight each response ratio (coupled with the amount of heterogeneity, i.e. the variance in the underlying effects)^30^.

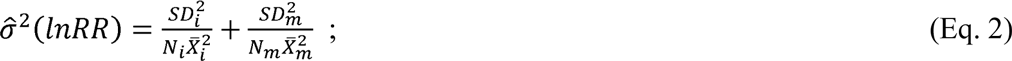

SDs were extracted from raw data when possible. If ranges were provided instead of SD (or SE or CV), we calculated SD following^613^. If neither ranges nor measures of variation were reported, but the reported sample size was > 1, we imputed SD based on the coefficient of variation from all complete cases (“Bracken approach”^614^). Imputation was done for 22% of all cases in mammals, 1.1% in birds, 11% in reptiles and 7.3% in amphibians, all within the upper limit of imputations (<30% of all cases per group) advised in previous studies^90^.

For each study and island-mainland comparison, we compiled the mainland and island names, the study reference, the body size index used, the geographic coordinates, the distance to the closest mainland (spatial isolation, km) and the island area (km^2^). We completed missing data on island characteristics using the UNEP island database (http://islands.unep.ch/) and the Threatened Island Biodiversity Database (TIB, http://tib.islandconservation.org/). Missing information was calculated using Google Earth. Additionally, we extracted the Normalized Difference Vegetation Index (NDVI) as a proxy for resource availability on islands^615^. We also calculated the standard deviation of NDVI to assess seasonality in leaf or vegetation cover, as an index of seasonality in available resources. NDVI was downloaded from NASA Ames Ecological Forecasting Lab (https://lpdaacsvc.cr.usgs.gov/appeears/task/area).

Because climate influences both resource requirements and primary productivity, body size evolution should also be influenced by climatic conditions on islands. We thus extracted island climatic conditions from WorldClim v. 2.0 (http://worldclim.org616). Specifically, we used variables that are more closely associated with the proposed underlying mechanisms of Bergmann’s rule (i.e. thermoregulation and starvation resistance): mean annual temperature, annual precipitation, and seasonality of temperature and precipitation^617^. We assumed that the time period for these bioclimatic variables (1970–2000), although not necessarily matching the actual time period of body size evolution in the insular populations, roughly represents the climatic conditions in the Holocene, a period relatively climatically stable where most of our populations became isolated (i.e., after the last glacial maximum; see also^9^). Because climatic variability across cells substantially exceeds variation within cells in the Holocene, current layers are considered adequate for geographic comparisons. All spatial variables were downloaded at 0.1-degree resolution, and we averaged all cells per island to obtain a mean value of each environmental variable (e.g., temperature, NDVI, precipitation, etc.). Finally, for each species included in our dataset, we collated diet information from EltonTraits for birds and mammals^618^, and from other sources for reptiles^608, 619^, and classified species as carnivores (> 50% diet consisting of vertebrates) or non-carnivores (< 50% diet consisting of vertebrates), following previous studies^79, 620^. As all amphibians in our dataset are carnivores^621^, we did not record their diet.

### Data analyses

To test the island rule hypothesis, we used phylogenetic meta-regressions between *lnRR* and body mass of mainland relatives, following most previous studies of the island rule (e.g. ^4, 5, 7^ ^6, 622, 623^). A negative slope for this relationship would support the island rule (Fig. 1).

The use of multiple populations of the same species can overestimate the actual number of degrees of freedom, generating type-1 errors. We controlled for this by adding ‘Species’ as a random effect intercept in our analyses. Additionally, body size evolution in insular vertebrates is heavily influenced by phylogenetic effects, with species within entire clades seemingly showing either dwarfism or gigantism^6^. Thus, we accounted for phylogeny by including the phylogenetic relatedness correlation matrix as a random effect. The species term captures the similarities of effect sizes within the same species, while the phylogenetic term represents the similarity due to relatedness^624^. We also added ‘Source’ as a random effect intercept to account for between-source variability and the fact that we had multiple response ratios per study. In some cases, ‘Source’ represented the combination of two sources of data, one for the island size and one for the mainland size. Finally, we included an observation level random effect, which represents the residual variance that needs to be explicitly modelled in a meta-analysis^29^. Total heterogeneity, and heterogeneity due to phylogeny, source and species identity were computed following Nakagawa & Santos (2012)^29^.

We tested the robustness of our results against several potential limitations. Because multiple island populations were often compared with a single mainland population, we accounted for these repeated measures in a variance-covariance matrix where the diagonal includes the sampling variances, and the off-diagonals of the matrix represent the shared variance (covariance) among the response ratios due to the common mainland population^625^. Further, we compared our main results to models fitted with *lnRR* and sampling variances corrected for small sample size^38^. Another potential problem is that regressions using ratios may lead to spurious correlations^31, 32^. Thus, we conducted an additional analysis testing the statistical significance of body size trends by regressing island mass against mainland mass, following previous studies^4, 5, 20, 41^. Phylogenetic meta-regressions were run using island mass as the response variable, and mainland mass as the predictor (both transformed with natural-log), with random effects as specified above, and sampling variance sd_i_ /mass_i_ *N_i_. This approach has some limitations in being harder to visualize and less effective in considering the sampling variance of measurements (representing intrapopulation variability), yet nonetheless provides an alternative approach for assessing the robustness of our results, in line with previous studies^4, 5, 20, 41^. Finally, we assessed publication bias by testing the influence of data source on the relationship between size ratio and mainland mass. This involved comparing whether patterns differed in island-mainland pairs extracted from studies testing the island rule (38.6% of cases) versus pairs extracted from studies not testing the island rule (61.4% of cases).

### Testing ecological hypotheses explaining insular size shifts

To evaluate the relative role of key mechanisms proposed to influence body size evolution in island fauna, we compiled a further range of variables (Supplementary Table 1, Extended Data Fig. 1). These included island area (linked to both resource limitation and to ecological release from both predation and competition) and spatial isolation (linked to reduced colonization from mainland populations for smaller taxa, i.e. immigration selection^50^). In addition, we included climatic and resource seasonality, which are linked to the starvation resistance hypothesis, and productivity and species diet, each of which are linked to resource limitation. Because body size evolution may be influenced by climate (e.g. Bergmann’s rule)^9, 626^, we also included mean temperature, which is linked to body size adaptations for enhancing heat conservation or dissipation (thermoregulation hypothesis). For amphibians, we included precipitation as a proxy for water supply linked to aquatic habitats, moisture and humidity (water availability hypothesis).

We modelled interactions between body size and each of the explanatory variables because we expected these factors to differentially affect species of different sizes, thus producing different effects in small, medium-sized and large species. In line with the ecological release and resource limitation hypotheses, we expected the slope of the *lnRR*-mainland mass relationship to be steeper in smaller islands, isolated from the mainland and with fewer or no predators (Fig. 1). Further, if resource availability is a key factor, we also expected large species to undergo dwarfism on islands with low productivity^48, 49^, and for dwarfism to be accentuated in dietary niches with high energy requirements, including carnivory^9^. In addition, high seasonality in resources and in temperature was expected to result in increased gigantism in small-sized species, because energy reserves increase faster than energy depletion as body size increases (starvation resistance hypothesis)^9, 627^. We hypothesized that smaller species would benefit comparatively more by increasing in size than larger species. Because amphibians are generally small-sized, we also fitted a model for this group with only additive terms (mainland mass + sdNDVI) where seasonality in resources would result in larger body sizes for all species. Finally, mechanisms driven by thermoregulation and water availability predict that body size shifts are associated with temperature and rainfall, respectively. Mean temperature was expected to predominantly affect endotherms and small ectotherms with good thermoregulating abilities (reptiles and anurans) living on cold islands which, compared to similar-sized species on islands with a mild climate, would exhibit more pronounced gigantism to enhance heat conservation. We fitted the effect of temperature as an interactive (mainland mass × Tmean) or additive term (mainland mass + Tmean) to assess whether only small species or all species would increase in size in low temperature islands (see details in Supplementary Table 1, Extended Data Fig 1, Supplementary Table 7).

Prior to modeling, all the moderators (explanatory variables) were inspected and log_10_-transformed if necessary to meet normality assumptions in model errors. We considered a result to be significant when the 95% confidence interval (CI) did not cross zero. We assessed the explained heterogeneity using Omnibus test for moderators (*Q_m_*) and the percentage of variance explained by the moderators using *R^2^* marginal^628^. All figures show the relationship between size response ratio and body mass, and how this might be altered by the mechanisms explained above.

All analyses were performed in R 3.5.3^629^ using the packages *metafor v2.0*^630^ and *metagear v0.4*^631^ for the meta-regression models and data imputation, *metaDigitise v1.0*^632^ for data extraction from plots, *ape v5.2*^633^ for estimating branch lengths and resolving polytomies, *rotl v3.0.4*^634^ for building the phylogenies for our species by searching the Open Tree Taxonomy^635^ and retrieving the phylogenetic relationships from the Open Tree of Life^636^, *sf v0.7-3* ^637^ and *raster v2.7-15*^638^ for spatial analyses, *dplyr v0.8.0.1*^639^ and *reshape2 v1.4.3*^640^ for data manipulation and *ggplot2 v 3.3.0.9000*^641^ and *ggpubr v0.1.8* ^642^ for data visualization. ArcMap 10.5 was used for Figure 2. Silhouettes in figures were extracted from ‘phylopic’ (https.phylopic.org). The PRISMA Checklist for systematic reviews is available in Appendix 3.

## Data availability

All data are available at https://github.com/anabenlop/Island_Rule and https://figshare.com/projects/Body_size_evolution_in_insular_vertebrates/89102.

## Code availability

The code to conduct the analyses is available at https://github.com/anabenlop/Island_Rule.

## Acknowledgements

We are grateful to K. B. Aubry, J. E. Keehn, S. Michaelides and D. Strickland for sharing their data with us, S. Meiri for pointing out valuable sources of measurement data, and to A. Sánchez-Tojar, P. Peres-Neto and S. Nakagawa for useful discussion on the analytical framework. ABL was supported by a Juan de la Cierva-Incorporación grant (IJCI-2017-31419) from the Spanish Ministry of Science, Innovation and Universities. LS and MAJH were supported by the ERC project (62002139 ERC – CoG SIZE 647224). We thank numerous biological collections, in particular the Natural History Museum, Tring, for providing access to specimens. Bird trait data collection was supported by Natural Environment Research Council grant nos. NE/I028068/1 and NE/P004512/1 (to JAT).

## Author Contributions

AB-L conceived and coordinated the research, led the analyses and wrote the first draft; AB-L, LS, JG-Z, MAJH and JAT helped to develop the conceptual framework; LS compiled the environmental rasters; JAT, PW and BM provided morphometric data. All authors contributed to the data collection from the literature and to the writing of the final manuscript.

