## Extended Data Figures for "The island rule explains consistent patterns of body size evolution in terrestrial vertebrates"

Extended Data for “The island rule explains consistent patterns of body size evolution across terrestrial vertebrates”

### **Extended Data Fig. 1. Conceptual models depicting the different hypotheses tested to explain insular size shifts in vertebrates.**

### **Extended Data Fig. 2.** Phylogenetic meta-regression models of mean body size (ln-transformed mass, in g) on islands versus mainland for (a) mammals, (b) birds, (c) reptiles and (d) amphibians. The dashed line has a slope of 1 and an intercept of 0. The solid lines represent the phylogenetic meta-regression slope estimate. The intercept a and slope b are presented in each plot along with CI. The island rule holds if the intercept is > 0 and the slope < 1.

### **Extended Data Fig. 3. Variation accounted for by random factors (Source, Species and Phylogeny) and residual variation. The amount of variance accounted for by phylogeny was the largest for reptiles and mammals, and low for birds. The extent of variance explained by data sources was larger for mammals and reptiles, and low for amphibians and birds. The residual variance was highest for birds, followed by mammals, amphibians and reptiles, indicating that other factors besides mainland body size may help to explain insular size shifts (see Extended Data Fig. 5-8).**

### **Extended Data Fig. 4. Phylogenetic random effects across taxonomic orders for (a) mammals and (b) birds; and across families for (c) reptiles and (d) amphibians.**

### **Extended Data Fig. 5. Ecological factors explaining insular size shifts in mammals. Only 1-2 variables are displayed per plot, while keeping the other predictors at median values. Continuous variables are represented at the 10% and 90% quantile for each extreme (close vs remote or small vs large, and low vs high). *lnRR* > 0 indicates gigantism, *lnRR* < 0 indicates dwarfism, and *lnRR* = 0 indicates no shift in body size in islands compared to mainland populations. Shaded areas represent 95% CI. QM indicates the explained heterogeneity (variance) by the interaction between each explanatory factor and body mass (e.g. mass:area in panel a), or the explanatory factor only in case of intercept-only models (e.g. temperature in this case). See Supplementary Table 7 for details.**

### **Extended Data Fig. 6. Ecological factors explaining insular size shifts in birds. Only 1-2 variables are displayed per plot, while keeping the other predictors at median values. Continuous variables are represented at the 10% and 90% quantile for each extreme (close vs remote or small vs large, and low vs high). *lnRR* > 0 indicates gigantism*, lnRR* < 0 indicates dwarfism, and *lnRR* = 0 indicates no shift in body size in islands compared to mainland populations. Shaded areas represent 95% CI. QM indicates the explained heterogeneity (variance) by the interaction between each explanatory factor and body mass (e.g. mass:area in panel a), or the explanatory factor only in case of intercept-only models (e.g. temperature in this case). See Supplementary Table 7 for details.**

### **Extended Data Fig. 7. Ecological factors explaining insular size shifts in reptiles. Only 1-2 variables are displayed per plot, while keeping the other predictors at median values. Continuous variables are represented at the 10% and 90% quantile for each extreme (close vs remote or small vs large, and low vs high). *lnRR* > 0 indicates gigantism, *lnRR* < 0 indicates dwarfism, and *lnRR* = 0 indicates no shift in body size in islands compared to mainland populations. Shaded areas represent 95% CI. QM indicates the explained heterogeneity (variance) by the interaction between each explanatory factor and body mass (e.g. mass:area in panel a), or the explanatory factor only in case of intercept-only models (e.g. temperature in this case). See Supplementary Table 7 for details.**

### **Extended Data Fig. 8. Ecological factors explaining insular size shifts in amphibians. Only 1-2 variables are displayed per plot, while keeping the other predictors at median values. Continuous variables are represented at the 10% and 90% quantile for each extreme (close vs remote or small vs large, and low vs high). *lnRR* > 0 indicates gigantism, *lnRR* < 0 indicates dwarfism, and *lnRR* = 0 indicates no shift in body size in islands compared to mainland populations. Shaded areas represent 95% CI. QM indicates the explained heterogeneity (variance) by the interaction between each explanatory factor and body mass (e.g. mass:area in panel a), or the explanatory factor only in case of intercept-only models (e.g. seasonality in resources – sdNDVI in this case). See Supplementary Table 7 for details.**
