## Supplementary Information for "The island rule explains consistent patterns of body size evolution in terrestrial vertebrates"

This document contains:

1. Supplementary Tables 1-8.
2. Supplementary Figure 1. Location of insular populations included in our analyses, with each panel representing a taxonomic group.
3. Supplementary Figure 2. The generality of the island rule in birds when tarsus length was used preferably to convert to body mass equivalents using allometric relationships.
4. Supplementary References.
5. Appendix 1. Databases searched within WOS.
6. Appendix 2. Structured Summary.
7. Appendix 3. PRISMA Checklist.

As separate Excel files:

Supplementary Dataset 1: Database of insular-mainland size ratios, species identity, island and mainland localities, sampling variance, and physiographic, climatic and ecological characteristics of islands. Located at https://github.com/anabenlop/Island_Rule.

Supplementary Dataset 2: Parameter estimates of the phylogenetic meta–regression models testing different ecological hypotheses that may explain body size evolution in islands. Located at https://github.com/anabenlop/Island_Rule

Supplementary Dataset 3:  List of exluded references and reasons for exclusion. Located at https://github.com/anabenlop/Island_Rule

Supplementary Dataset 4: Skull, length and weight data used to fit allometric relationships that were not available in the literature. Located at https://github.com/anabenlop/Island_Rule

**Supplementary Table 1. Main factors that may influence body size evolution of terrestrial vertebrates (mammals, birds, reptiles and amphibians) in islands.**

| Predictor | Ecological hypothesis | Rationale | Taxonomic groups |
|---|---|---|---|
| Mass of mainland ancestor/sister species | Island rule *sensu stricto* | Insular body size shifts vary among differently sized animals: Small species are predicted to become larger (gigantism) whereas large animals will become smaller (dwarfism). For amphibians, most studies show an overall tendency to gigantism because they are usually small and they are the principal prey of other predators[1]. | All |
| Island area ($km^2$) | Ecological release hypothesis/Resource limitation hypothesis | Small islands are usually less productive and thus becoming smaller is adaptive for large species with high energetic requirements[2]. Smaller species would become larger because few/no predators and large competitors inhabit small islands resulting in greater utilization of the island resources and intraspecific competition. Large species would become smaller when released from predation pressure (e.g., ungulates or lagomorphs in islands with no large carnivores)[2], because investing energy in size is no longer adaptive as a defense from predators, thereby reallocating energy to other functions such as reproduction. | All |
| Distance to mainland (spatial isolation, km) | Immigration selection hypothesis | Immigration rate and gene flow are limited in isolated islands. More isolated islands should be colonized by larger species because larger animals should have greater physiological endurance and dispersal capacities. Thus, body size of insular populations of a particular species should increase with island isolation. This affects mostly small species with limited dispersal capacities will not reach isolated islands, as only the largest individuals in the population will be able to colonize more isolated islands, leading to a stronger signal in insular size shifts[2]. In turn, in islands close to the mainland (i.e.: land bridge, continental islands), we expect that gene flow has not been impaired, either because many of these islands were connected to the continent in quite recent times (i.e. Last Glacial Maximum), and there has been insufficient time for differences to accumulate; or because several recolonization events have occurred in both directions, leading to hybridization between insular and mainland relatives. | Small species in mammals and reptiles. All amphibians. |
| Average temperature | Thermoregulation hypothesis | The island rule will be exacerbated in cold islands for small endotherm species, which are expected to become larger to increase their surface area to volume ratio and thus lose less heat relative to their mass. In turn, to improve heat dissipation, in warm islands large endotherms would tend even more towards dwarfism than predicted by the island rule only[2,3]. In ectotherms with good thermoregulating abilities (reptiles and anurans), small species will tend to larger sizes in colder islands than expected by the island rule only because of enhanced heat retention[4]. Alternatively, all species regardless of the ancestral body size will be larger in colder islands, and smaller in warmer islands, with changes in the intercept of the size ratio-mainland mass relationship, but not in the slope. | Small and large endotherms and small ectotherms |
| Precipitation | Water availability hypothesis | We expect that small amphibian species are larger in islands with low precipitation than predicted by the island rule only because larger body size in amphibians is adaptive in drier environments due to a lower surface/mass ratio that reduces the loss of water[5,6]. | Amphibians |
| Resource availability (NDVI) | Resource limitation hypothesis | Since resource requirements tend to increase with body size, less productive islands are expected to exert a stronger selective pressure toward smaller sizes in large-bodied species[2,7,8]. | Large species across all groups |
| Seasonality in temperature (Tseas) or in resources (NDVIsd) | Starvation resistance hypothesis | Islands with high temperature seasonality or high seasonality in resources may favor individuals of larger size in small species because energy reserves increase faster than energy depletion as body size increases, which is adaptive in seasonal environments where animals experience long periods without food (e.g. during aestivation or hibernation)[2,9]. | Small species across all groups. All amphibians. |
| Diet (carnivores, non-carnivores) | Resource limitation hypothesis | Carnivores are expected to respond more clearly to selective pressures on islands due to their higher energetic requirements[2,10]. | Mammals<br>Birds<br>Reptiles |

**Supplementary Table 2. Allometric relationships used to convert body size indexes to mass. CBL: condylobasal length, SCL: Straight carapace length, SVL: Snout-vent length, TL: Total length. All relationships are based on OLS (Ordinary Least Square) models except those by Santini et al. 2018 which used PGLS (Phylogenetic Generalized Least Square) models. NR: Not reported. For some relationships the underlying data was completed with data from different sources (Supplementary Dataset 3)**

| Class | Order | Family | Species | Measure | Equation | N | $R^2$ | Source |
|---|---|---|---|---|---|---|---|---|
| Amphibians | Anura | All | All | SVL | $\log_{10}$ (mass, g) = -4.324 + 3.189 $\log_{10}$ (SVL, mm) | 88 | 0.950 | Santini, et al. [11] |
| Amphibians | Anura | Bufonidae | All | SVL | $\log_{10}$ (mass, g) = -3.791 + 2.914 $\log_{10}$ (SVL, mm) | 9 | 0.980 | Santini, et al. [11] |
| Amphibians | Anura | Dicroglossidae | Fejervarya limnocharis | SVL | $\log_{10}$ (mass, g) = -4.431 + 3.209 $\log_{10}$ (SVL, mm) | NR | 0.963 | Thammachoti, et al. [12] |
| Amphibians | Anura | Hylidae | All | SVL | $\log_{10}$ (mass, g) = -4.462 + 3.201 $\log_{10}$ (SVL, mm) | 35 | 0.938 | Santini, et al. [11] |
| Amphibians | Anura | Myobatrachidae | All | SVL | $\log_{10}$ (mass, g) = -4.586 + 3.372 $\log_{10}$ (SVL, mm) | 12 | 0.952 | Santini, et al. [11] |
| Amphibians | Anura | Ranidae | All | SVL | $\log_{10}$ (mass, g) = -4.862 + 3.492 $\log_{10}$ (SVL, mm) | 13 | 0.847 | Santini, et al. [11] |
| Amphibians | Caudata | Plethodontidae | All | SVL | $\log_{10}$ (mass, g) = -4.706 + 2.968 $\log_{10}$ (SVL, mm) | 21 | 0.925 | Santini, et al. [11] |
| Amphibians | Caudata | Salamandridae | All | SVL | $\log_{10}$ (mass, g) = -4.744 + 3.073 $\log_{10}$ (SVL, mm) | 13 | 0.933 | Santini, et al. [11] |
| Amphibians | Gymnophiona | All | All | TL | $\log_{10}$ (mass, g) = -5.465 + 2.597 $\log_{10}$ (SVL, mm). | | | Feldman, et al. [13]. Allometry for snakes used as approximation (see also Pough 1980) |
| Birds | All | All | All | Bill length | $\log_{10}$ (mass, g) = -0.868 + 1.922 $\log_{10}$ (bill, mm) | 2376 | 0.530 | Lislevand, et al. [14] |
| Birds | All | All | All | Tarsus length | $\log_{10}$ (mass, g) = -1.778 + 2.482 $\log_{10}$ (tarsus, mm) | 2257 | 0.690 | Lislevand, et al. [14] |
| Birds | All | All | All | Wing length | $\log_{10}$ (mass, g) = -3.399 + 2.508 $\log_{10}$ (wing, mm) | 2618 | 0.890 | Lislevand, et al. [14] |
| Mammals | Rodentia | Muridae | Apodemus argenteus | antero-posterior diameter of the lower incisor | ln (mass,g) = 3.19 + 2.58 ln (anteposterior incisor, mm) | 51 | 0.930 | Millien-Parra [15] |
| Mammals | All | All | All | CBL | $\log_{10}$ (mass, g) = -3.177 + 3.375 $\log_{10}$ (CBL, mm) | 357 | 0.971 | Supplementary Dataset 4, includes data from Van Valkenburgh [16] |
| Mammals | Carnivora | All | All | CBL | $\log_{10}$ (mass, g) = -3.307 + 3.439 $\log_{10}$ (CBL, mm) | 243 | 0.941 | Supplementary Dataset 4, includes data from Van Valkenburgh [16] |
| Mammals | Carnivora | Canidae | All | CBL | $\log_{10}$ (mass, kg) = -5.180 + 2.848 $\log_{10}$ (CBL, mm) | 14 | 0.883 | Van Valkenburgh [16] |
| Mammals | Carnivora | Felidae | All | CBL | $\log_{10}$ (mass, kg) = -5.592 + 3.170 $\log_{10}$ (CBL, mm) | 16 | 0.865 | Van Valkenburgh [16] |

| | | | | | | | | |
|---|---|---|---|---|---|---|---|---|
| Mammals | Carnivora | Mustelidae | All | CBL | $\log_{10}$ (mass, g) = -4.299 + 3.972 $\log_{10}$ (CBL, mm) | 75 | 0.956 | Supplementary Dataset 4 |
| Mammals | Carnivora | Ursidae | Ursus arctos | CBL | $\log_{10}$ (mass, g) = -3.999 + 3.615 $\log_{10}$ (CBL, mm) | 19 | 0.954 | Stringham [17] |
| Mammals | Carnivora | Herpestidae | All | CBL | $\log_{10}$ (mass, g) = -4.299 + 3.972 $\log_{10}$ (CBL, mm) | 75 | 0.956 | Supplementary Dataset 4 |
| Mammals | Carnivora | Viverridae | All | CBL | $\log_{10}$ (mass, g) = -4.299 + 3.972 $\log_{10}$ (CBL, mm) | 75 | 0.956 | Supplementary Dataset 4 |
| Mammals | Chiroptera | All | All | CBL | $\log_{10}$ (mass, g) = -1.706 + 2.227 $\log_{10}$ (CBL, mm) | 41 | 0.780 | Supplementary Dataset 4 |
| Mammals | Eulipotyphla | All | All | CBL | $\log_{10}$ (mass, g) = -4.258 + 4.048 $\log_{10}$ (CBL, mm) | 18 | 0.990 | Supplementary Dataset 4 |
| Mammals | Rodentia | All | All | CBL | $\log_{10}$ (mass, g) = -3.753 + 3.718 $\log_{10}$ (CBL, mm) | 28 | 0.991 | Supplementary Dataset 4 |
| Mammals | Rodentia | Castoridae | All | CBL | $\log_{10}$ (mass, kg) = -5.245 +3.116 $\log_{10}$ (CBL, mm) | 76 | 0.952 | Reynolds [18] |
| Mammals | All | All | All | Length | $\log_{10}$ (mass, g) = -4.151 + 2.841 $\log_{10}$ (length, mm) | 3542 | 0.977 | Jones, et al. [19] |
| Mammals | Chiroptera | All | All | Length | $\log_{10}$ (mass, g) = -3.144 + 2.409 $\log_{10}$ (length, mm) | 350 | 0.880 | Nowak and Walker [20] |
| Mammals | Rodentia | All | All | Length | $\log_{10}$ (mass, g) = -4.100 + 2.821 $\log_{10}$ (length, mm) | 210 | 0.935 | Nowak and Walker [20] |
| Mammals | All | All | All | Skull | log10 (mass, g) = -2.872 + 3.165 $\log_{10}$ (Skull, mm) | 64 | 0.980 | Supplementary Dataset 4 |
| Mammals | Carnivora | All | All | Skull | $\log_{10}$ (mass, kg) = -7.767 + 4.024 $\log_{10}$ (Skull, mm) | 39 | 0.976 | Figueirido, et al. [21] |
| Mammals | Cetartiodactyla | All | All | Skull | $\log_{10}$ (mass, g) = -2.821 + 3.126 $\log_{10}$ (Skull, mm) | 25 | 0.946 | Fitch [22] |
| Mammals | Lagomorpha | All | All | Skull | $\log_{10}$ (mass, g) = -2.999 + 3.285 $\log_{10}$ (Skull, mm) | 17 | 0.917 | Kraatz, et al. [23] |
| Mammals | Primata | All | All | Skull | $\log_{10}$ (mass, g) = -2.463 + 3.071 $\log_{10}$ (Skull, mm) | 90 | 0.916 | Plavcan and Ruff [24] |
| Reptiles | Testudines | All | All | SCL | $\log_{10}$ (mass, g) = -3.855 + 2.677 $\log_{10}$ (SCL, mm) | 692 | 0.951 | Regis and Meik [25] |
| Reptiles | Squamata | Agamidae | All | SVL | $\log_{10}$ (mass, g) = -4.686 + 3.105 $\log_{10}$ (SVL, mm) | 83 | 0.965 | Feldman, et al. [13] |
| Reptiles | Squamata | Anguidae | All | SVL | $\log_{10}$ (mass, g) = -5.765 + 3.480 $\log_{10}$ (SVL, mm) | 11 | 0.897 | Feldman, et al. [13,26] |
| Reptiles | Squamata | Boidae | All | SVL | $\log_{10}$ (mass, g) = -5.500 + 2.776 $\log_{10}$ (SVL, mm) | 15 | 0.883 | Feldman and Meiri [27] |
| Reptiles | Squamata | Chamaeleonidae | All | SVL | $\log_{10}$ (mass, g) = -3.997 + 2.680 $\log_{10}$ (SVL, mm) | 23 | 0.970 | Meiri [28] |
| Reptiles | Squamata | Colubridae | All | SVL | $\log_{10}$ (mass, g) = -5.525 + 2.628 $\log_{10}$ (SVL, mm) | NR | NR | Feldman, et al. [13], Feldman and Meiri [27] |
| Reptiles | Squamata | Dactyloidae | All | SVL | $\log_{10}$ (mass, g) = -4.574 + 2.942 $\log_{10}$ (SVL, mm) | 95 | 0.930 | Novosolov, et al. [29] |
| Reptiles | Squamata | Dibamidae | All | SVL | $\log_{10}$ (mass, g) = -4.207 + 2.300 $\log_{10}$ (SVL, mm) | 24 | 0.758 | Meiri [28] |
| Reptiles | Squamata | Diplodactylidae | All | SVL | $\log_{10}$ (mass, g) = -4.804 + 3.057 $\log_{10}$ (SVL, mm) | 39 | 0.900 | Scharf, et al. [30] |
| Reptiles | Squamata | Dipsadidae | All | SVL | log10 (mass, g) = -5.219 + 2.561 log10 (SVL, mm) | NR | NR | Feldman, et al. [13] |
| Reptiles | Squamata | Elapidae | All | SVL | log10 (mass, g) = -4.892 + 2.453 log10 (SVL, mm) | 26 | 0.844 | Feldman and Meiri [27] |

| | | | | | | | | |
|---|---|---|---|---|---|---|---|---|
| Reptiles | Squamata | Gekkonidae | All | SVL | log10 (mass, g) = -4.242 + 2.761 log10 (SVL, mm) | 66 | 0.850 | Novosolov, et al. [29] |
| Reptiles | Squamata | Gymnophthalmidae | All | SVL | log10 (mass, g) = -5.178 + 3.302 log10 (SVL, mm) | 29 | 0.804 | Meiri [28] |
| Reptiles | Squamata | Iguanidae | All | SVL | log10 (mass, g) = -4.298 + 2.972 log10 (SVL, mm) | 24 | 0.895 | Meiri [28] |
| Reptiles | Squamata | Lacertidae | All | SVL | log10 (mass, g) = -4.543 + 2.951 log10 (SVL, mm) | 87 | 0.881 | Meiri [28] |
| Reptiles | Squamata | Lamprophiidae | All | SVL | log10 (mass, g) = -7.092 + 3.232 log10 (SVL, mm) | 35 | 0.886 | Feldman and Meiri [27] |
| Reptiles | Squamata | Natricidae | All | SVL | log10 (mass, g) = -6.223 + 2.982 log10 (SVL, mm) | NR | NR | Scharf, et al. [30] |
| Reptiles | Squamata | Phrynosomatidae | All | SVL | log10 (mass, g) = -3.855 + 2.677 log10 (SVL, mm) | 40 | 0.852 | Meiri [28] |
| Reptiles | Squamata | Phyllodactylidae | All | SVL | log10 (mass, g) = -4.482 + 2.945 log10 (SVL, mm) | 17 | 0.967 | Scharf, et al. [30] |
| Reptiles | Squamata | Scincidae | All | SVL | log10 (mass, g) = -5.125 + 3.229 log10 (SVL, mm) | 154 | 0.957 | Meiri [28] |
| Reptiles | Squamata | Sphaerodactylidae | All | SVL | log10 (mass, g) = -4.559 + 2.970 log10 (SVL, mm) | 24 | 0.960 | Novosolov, et al. [29] |
| Reptiles | Squamata | Teiidae | All | SVL | log10 (mass, g) = -4.747 + 3.110 log10 (SVL, mm) | 43 | 0.960 | Meiri [28] |
| Reptiles | Squamata | Varanidae | All | SVL | log10 (mass, g) = -5.301 + 3.235 log10 (SVL, mm) | 45 | 0.960 | Meiri [28] |
| Reptiles | Squamata | Viperidae | All | SVL | log10 (mass, g) = -5.165 + 2.655 log10 (SVL, mm) | 60 | 0.913 | Feldman and Meiri [27] |
| Reptiles | Squamata | Xantusiidae | All | SVL | log10 (mass, g) = -4.796 + 3.048 log10 (SVL, mm) | 7 | 0.940 | Meiri [28] |
| Reptiles | Squamata | Boidae | All | TL | log10 (mass, g) = -5.886 + 2.856 log10 (TL, mm) | 13 | 0.871 | Feldman and Meiri [27] |
| Reptiles | Squamata | Colubridae | All | TL | log10 (mass, g) = -5.548 + 2.539 log10 (TL, mm) | 154 | 0.792 | Feldman and Meiri [27] |
| Reptiles | Squamata | Dipsadidae | All | TL | log10 (mass, g) = -4.715 + 2.278 log10 (TL, mm) | 69 | 0.680 | Scharf, et al. [30] |
| Reptiles | Squamata | Lamprophiidae | All | TL | log10 (mass, g) = -6.286 + 2.821 log10 (TL, mm) | 31 | 0.839 | Feldman and Meiri [27] |
| Reptiles | Squamata | Viperidae | All | TL | log10 (mass, g) = -6.103 + 2.910 log10 (TL, mm) | 51 | 0.877 | Feldman and Meiri [27] |

**Supplementary Table 3. Parameter estimates for the phylogenetic meta-regression models testing the generality of the island rule in terrestrial vertebrates, using $lnRR^{\Delta}$ and sampling variance corrected for small sample size[31]. *k:* number of island-mainland comparisons ($lnRR^{\Delta}$), $Q_m$: test of moderators (log$_{10}$(mainland mass). $R^2_m$: marginal $R^2$, estimated percentage of heterogeneity explained by the moderator (fixed effects). $R^2_c$: conditional $R^2$, percentage of heterogeneity attributable to fixed and random effects.**

| Class | *k* | Intercept (CI) | Slope (CI) | $Q_m$ (p-value) | $R^2_m$ | $R^2_c$ |
|---|---|---|---|---|---|---|
| Mammals | 1058 | 0.208 (0.053 – 0.364) | -0.088 (-0.121 – -0.055) | 27.31 (p < 0.001) | 11.4 | 56.1 |
| Birds | 695 | 0.216 (0.117 – 0.315) | -0.104 (-0.145 – -0.064) | 25.41 (p < 0.001) | 7.0 | 43.4 |
| Reptiles | 547 | 0.410 (0.006 – 0.813) | -0.305 (-0.419 – -0.190) | 27.24 (p < 0.001) | 17.7 | 66.6 |
| Amphibians | 179 | 0.195 (0.012 – 0.377) | -0.107 (-0.320 – 0.107) | 0.95 (p = 0.329) | 1.4 | 67.1 |

**Supplementary Table 4. Parameter estimates of the relationship between ln(island mass) and ln(mainland mass) as an alternative, complementary approach to our modelling framework. Models with intercept > 0 and slope < 1 would support the island rule (see also Lomolino 1985, 2005[1,32] and Meiri et al. 2011[33] for similar approaches regressing island size against mainland size). test: either Z value to test H0 intercept = 0, or t-value for the H0 slope = 1.**

| Class | Variable | Estimate | Lower 95CI | Upper 95 CI | test | P(test) |
|---|---|---|---|---|---|---|
| Mammals | Intercept | 0.208 | 0.049 | 0.367 | 2.57 | 0.010 |
| | ln(mainland mass) | 0.964 | 0.951 | 0.976 | 5.19 | <0.001 |
| Birds | Intercept | 0.215 | 0.115 | 0.314 | 4.22 | <0.001 |
| | ln(mainland mass) | 0.955 | 0.937 | 0.972 | 5.01 | <0.001 |
| Reptiles | Intercept | 0.410 | 0.002 | 0.818 | 1.97 | 0.049 |
| | ln(mainland mass) | 0.868 | 0.818 | 0.918 | 5.19 | <0.001 |
| Amphibians | Intercept | 0.194 | 0.012 | 0.375 | 2.09 | 0.036 |
| | ln(mainland mass) | 0.955 | 0.862 | 1.047 | 1.12 | 0.962 |

**Supplementary Table 5. Parameter estimates of the relationship between body size divergence and mainland mass for island-mainland comparisons where SDs were reported by the authors (i.e. no imputation).** ***k:*** **number of island-mainland comparisons (*lnRR)*, $Q_m$: test of moderators (log₁₀(mainland mass).** $R^2_m$: **marginal** $R^2$**, estimated percentage of heterogeneity explained by the moderator (fixed effects).** $R^2_c$: **conditional** $R^2$**, percentage of heterogeneity attributable to fixed and random effects.**

| Class | *k* | Intercept (CI) | Slope (CI) | $Q_m$ (p-value) | $R^2_m$ | $R^2_c$ |
|---|---|---|---|---|---|---|
| Mammals | 792 | 0.172 (0.052 – 0.292) | -0.083 (-0.116 – -0.051) | 25.38 ($p < 0.001$) | 10.7 | 55.4 |
| Birds | 681 | 0.214 (0.114 – 0.314) | -0.105 (-0.146 – -0.064) | 24.96 ($p < 0.001$) | 6.9 | 42.9 |
| Reptiles | 484 | 0.461 (0.002 – 0.923) | -0.343 (-0.463 – -0.224) | 31.71 ($p < 0.001$) | 20.5 | 66.5 |
| Amphibians | 166 | 0.194 (0.004 – 0.385) | -0.107 (-0.327 – 0.113) | 0.91 ($p = 0.340$) | 1.2 | 69.7 |

**Supplementary Table 6. Sensitivity analysis testing the relationship between body size divergence and mainland mass for studies testing or not the island rule (potential publication bias).** ***k:*** **total number of island-mainland comparisons, which is partitioned into the number of island-mainland comparisons from studies testing the island rule ($k_{ir}$) or not testing the island rule ($k_{nir}$).** ***lnRR:*** **body size divergence ratio,** $Q_m$***(full)*****: test of moderators for the full model.** $Q_m$***(data source)*****: test of moderators for the effect of Data source (studies testing the island rule vs studies not testing it).**

| Model | Class | *k* | $Q_{m\ (full)}$ (p-value) | $Q_{m\ (data\ source)}$ (p-value) |
|---|---|---|---|---|
| lnRR~ Mainland mass + Data source type | Mammals | 1058 ($k_{ir}$: 563, $k_{nir}$: 495) | 27.13 ($p < 0.001$) | 0.10 ($p = 0.746$) |
| | Birds | 695 ($k_{ir}$: 35, $k_{nir}$: 660) | 27.40 ($p < 0.001$) | 2.00 ($p = 0.157$) |
| | Reptiles | 547 ($k_{ir}$: 238, $k_{nir}$: 309) | 27.62 ($p < 0.001$) | 0.04 ($p = 0.834$) |
| | Amphibians | 179 ($k_{ir}$: 118, $k_{nir}$: 61) | 1.00 ($p = 0.606$) | < 0.01 ($p = 0.978$) |
| lnRR~ Mainland mass x Data source type | Mammals | 1058 ($k_{ir}$: 563, $k_{nir}$: 495) | 27.03 ($p < 0.001$) | 0.10 ($p = 0.756$) |
| | Birds | 695 ($k_{ir}$: 35, $k_{nir}$: 660) | 28.92 ($p < 0.001$) | 1.54 ($p = 0.214$) |
| | Reptiles | 547 ($k_{ir}$: 238, $k_{nir}$: 309) | 27.69 ($p < 0.001$) | 0.23 ($p = 0.633$) |
| | Amphibians | 179 ($k_{ir}$: 118, $k_{nir}$: 61) | 1.32 ($p = 0.724$) | 0.35 ($p = 0.555$) |

**Supplementary Table 7. Test of moderators for the phylogenetic meta-regression models testing different ecological hypotheses that may explain body size evolution in islands. *k:* number of island-mainland comparisons, *lnRR:* size divergence ratio, $Q_m$*(full)*: test of moderators for the full model. $Q_m$*(effect)*: test of moderators for the effect tested, which is the depicted in the inset figures (see also Extended Data Fig. 1). This could be an interaction effect where the slope of the lnRR-mainland mass relationship changes with varying values of the tested environmental or ecological factor, or an intercept effect, where the slope of the lnRR-mainland mass relationship does not change, but the intercept changes with varying values of the tested environmental or ecological factor. When two interactive terms are simultaneously tested (e.g.: mainland mass x distance + mainland mass x area), $Q_m$ *(effect)* accounts for the combined effect of both interactions. $R^2_{marginal}$ : estimated percentage of heterogeneity explained by the moderator. In bold those effects that were significantly different from zero (p < 0.05). Parameter estimates are available in Supplementary Dataset 2.**

| Hypothesis | Model | Class | *k* | $Q_m$ *(full)* (p-value) | $Q_m$ *(effect)* (p-value) | $R^2_m$ | Expected relationship |
|---|---|---|---|---|---|---|---|
| Ecological release<br>Resource limitation | lnRR~ Mainland mass x area | Mammals | 1058 | 34.03 (p < 0.001) | **5.85 (p = 0.016)** | 12.0 | 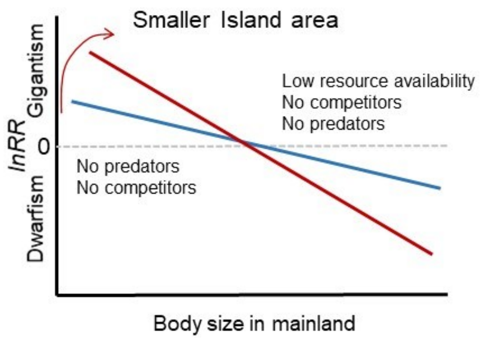 |
| | | Birds | 695 | 30.00 (p < 0.001) | 0.16 (p = 0.690) | 7.8 | |
| | | Reptiles | 547 | 38.68 (p < 0.001) | **6.85 (p = 0.009)** | 20.8 | |
| | | Amphibians | 179 | 5.12 (p = 0.164) | 3.05 (p = 0.081) | 4.2 | |
| Immigration selection | lnRR~ Mainland mass x distance | Mammals | 1058 | 34.83 (p < 0.001) | 3.55 (p = 0.06) | 11.6 | 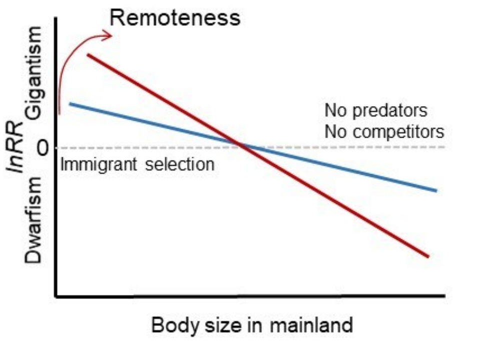 |
| | | Birds | 695 | 27.36 (p < 0.001) | 1.94 (p = 0.164) | 7.6 | |
| | | Reptiles | 547 | 30.02 (p < 0.001) | 2.02 (p = 0.155) | 17.5 | |
| | | Amphibians | 179 | 4.99 (p = 0.172) | 0.02 (p = 0.897) | 2.7 | |

| Hypothesis | Model | Class | *k* | $Q_m$ *(full)* **(p-value)** | $Q_m$ *(effect)* **(p-value)** | $R^2_m$ | Expected relationship |
|---|---|---|---|---|---|---|---|
| Ecological release<br>Resource limitation<br>Immigration selection | lnRR~ Mainland mass x area<br>+ Mainland mass x distance | Mammals<br><br>Birds<br><br>Reptiles<br><br>Amphibians | 1058<br><br>695<br><br>547<br><br>179 | 40.64<br>($p < 0.001$)<br>32.69<br>($p < 0.001$)<br>46.58<br>($p < 0.001$)<br>8.93<br>($p = 0.112$) | **12.20**<br>**($p = 0.002$)**<br>2.66<br>($p = 0.264$)<br>**12.18**<br>**($p = 0.002$)**<br>2.98<br>($p = 0.225$) | 12.6<br><br>8.5<br><br>20.7<br><br>5.1 | 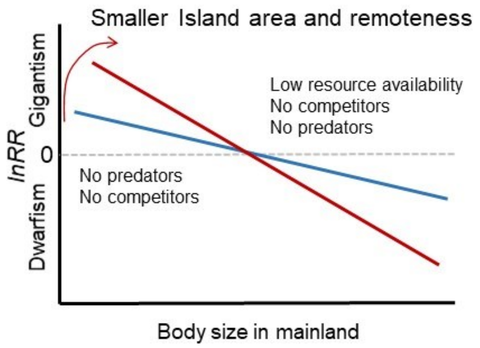 |
| Thermoregulation | lnRR~ Mainland mass x<br>tmean | Mammals<br><br>Birds<br><br>Reptiles<br><br>Amphibians | 1058<br><br>695<br><br>547<br><br>179 | 36.64<br>($p < 0.001$)<br>45.72<br>($p < 0.001$)<br>32.67<br>($p < 0.001$)<br>2.11<br>($p = 0.550$) | 0.12<br>($p = 0.726$)<br>**5.53**<br>**($p = 0.019$)**<br>1.45<br>($p = 0.229$)<br>1.09<br>($p = 0.297$) | 13.4<br><br>10.9<br><br>20.9<br><br>3.2 | 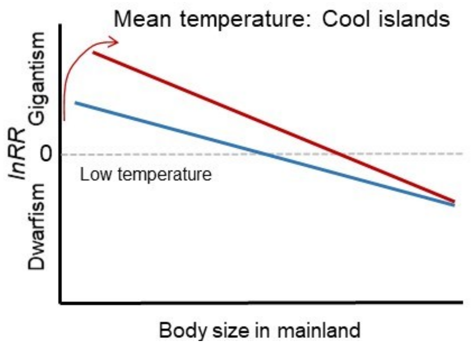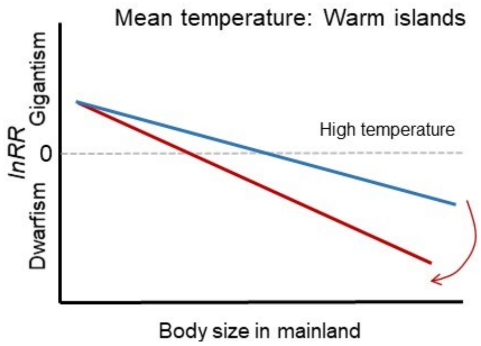 |

| Hypothesis | Model | Class | $k$ | $Q_{m\ (full)}$ (p-value) | $Q_{m\ (effect)}$ (p-value) | $R^2_m$ | Expected relationship |
|---|---|---|---|---|---|---|---|
| Thermoregulation | lnRR~ Mainland mass + tmean | Mammals | 1058 | 36.8 ($p < 0.001$) | **7.85 ($p = 0.005$)** | 13.7 | 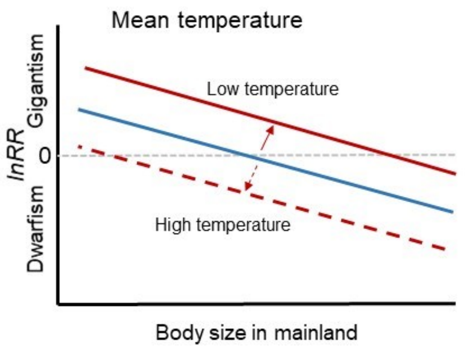 |
| | | Birds | 695 | 40.28 ($p < 0.001$) | **14.54 ($p < 0.001$)** | 10.2 | |
| | | Reptiles | 547 | 31.54 ($p < 0.001$) | **3.98 ($p = 0.046$)** | 18.7 | |
| | | Amphibians | 179 | 1. 02 ($p = 0.599$) | 0.008 ($p = 0.777$) | 1.4 | |
| Starvation resistance | lnRR~ Mainland mass x tseas | Mammals | 1058 | 27.99 ($p < 0.0001$) | 0.87 ($p = 0.351$) | 10.8 | 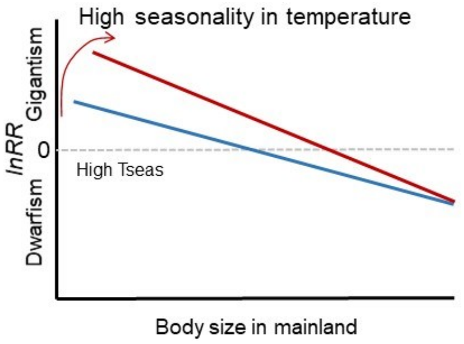 |
| | | Birds | 695 | 49.23 ($p < 0.001$) | **12.23 ($p < 0.001$)** | 10.9 | |
| | | Reptiles | 547 | 27.32 ($p < 0.001$) | 0.33 ($p = 0.566$) | 16.5 | |
| | | Amphibians | 179 | 2.25 ($p = 0.523$) | 0.66 ($p = 0.415$) | 4.3 | |
| Resource limitation | lnRR~ Mainland mass x NDVI | Mammals | 1058 | 30.03 ($p < 0.001$) | 1.02 ($p = 0.313$) | 11.3 | 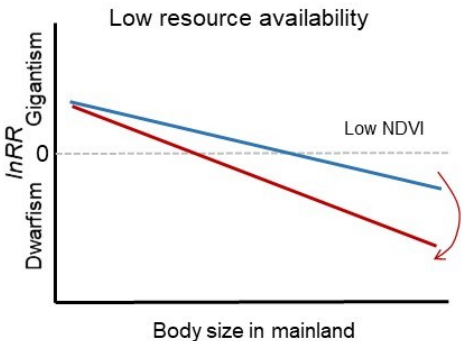 |
| | | Birds | 695 | 26.12 ($p < 0.001$) | 0.19 ($p = 0.662$) | 7.1 | |
| | | Reptiles | 547 | 34.41 ($p < 0.001$) | **5.74 ($p = 0.017$)** | 18.6 | |
| | | Amphibians | 179 | 0.99 ($p = 0.803$) | 0.01 ($p = 0.918$) | 1.4 | |

| Hypothesis | Model | Class | *k* | $Q_{m\ (full)}$ (p-value) | $Q_{m\ (effect)}$ (p-value) | $R^2_m$ | Expected relationship |
|---|---|---|---|---|---|---|---|
| Starvation resistance | lnRR~ Mainland mass x sdNDVI | Mammals | 1058 | 30.87 (p < 0.001) | 1.72 (p = 0.189) | 11.4 | 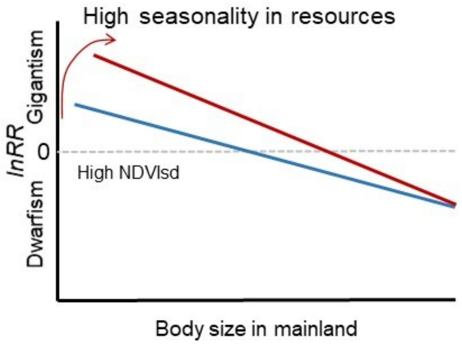 |
| | | Birds | 695 | 26.91 (p < 0.001) | 0.10 (p = 0.756) | 7.4 | |
| | | Reptiles | 547 | 34.64 (p < 0.001) | **6.83 (p = 0.009)** | 17.7 | |
| | | Amphibians | 179 | 11.92 (p = 0.008) | 0.358 (p = 0.550) | 8.6 | |
| Starvation resistance | lnRR~ Mainland mass + sdNDVI | Amphibians | 179 | 11.70 (p = 0.003) | **10.62 (p = 0.001)** | 8.1 | 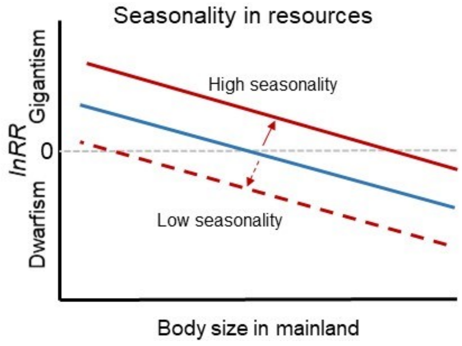 |
| Water availability | lnRR~ Mainland mass x prec | Amphibians | 179 | 1.30 (p = 0.728) | 0.17 (p = 0.679) | 1.6 | 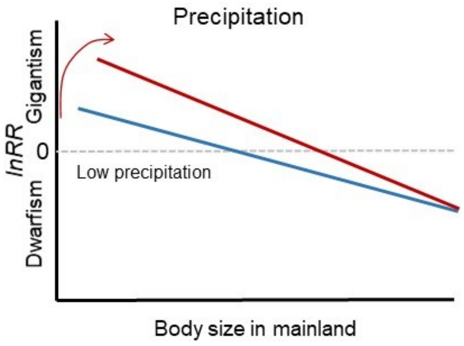 |

| **Hypothesis** | **Model** | **Class** | ***k*** | **$Q_{m\ (full)}$ (p-value)** | **$Q_{m\ (effect)}$ (p-value)** | **$R^2_m$** | **Expected relationship** |
|---|---|---|---|---|---|---|---|
| Resource limitation | lnRR~ Mainland mass x diet | Mammals<br><br>Birds<br><br>Reptiles | 1058<br><br>695<br><br>547 | 28.71<br>($p < 0.001$)<br>25.14<br>($p < 0.001$)<br>27.64<br>($p < 0.001$) | 0.19<br>($p = 0.664$)<br>0.01<br>($p = 0.927$)<br>0.61<br>($p = 0.435$) | 11.0<br><br>6.9<br><br>17.6 | 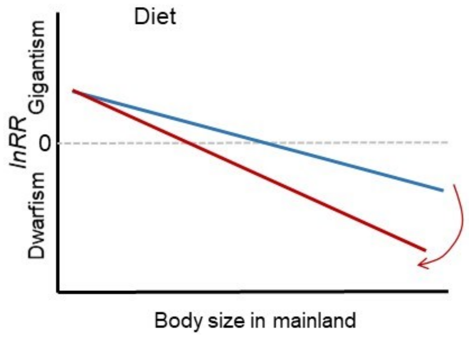 |

**Supplementary Table 8. Test of moderators for the phylogenetic meta-regression models testing the effects of island area and distance to mainland after removing archipelagos from the dataset. *k:* number of island-mainland comparisons, *lnRR:* size divergence ratio, $Q_m$*(full)*: test of moderators for the full model. $Q_m$*(effect)*: test of moderators for the effect tested. $R^2_{marginal}$: estimated percentage of heterogeneity explained by the moderator. In bold those effects that were significantly different from zero (p < 0.05). Parameter estimates are available in Supplementary Dataset 2.**

| Hypothesis | Model | Class | *k* | $Q_{m\ (full)}$ (p-value) | $Q_{m\ (effect)}$ (p-value) | $R^2_m$ |
|---|---|---|---|---|---|---|
| Ecological release<br>Resource limitation | lnRR~ Mainland mass x area | Mammals | 1038 | 32.91<br>(p < 0.001) | **6.22<br>(p = 0.016)** | 11.5 |
| | | Birds | 648 | 26.44<br>(p < 0.001) | 0.10<br>(p = 0.756) | 7.3 |
| | | Reptiles | 535 | 38.00<br>(p < 0.001) | **6.78<br>(p = 0.009)** | 20.6 |
| | | Amphibians | 179 | 5.12<br>(p = 0.164) | 3.05<br>(p = 0.081) | 4.8 |
| Immigration selection | lnRR~ Mainland mass x distance | Mammals | 1038 | 28.60<br>(p < 0.001) | 2.89<br>(p = 0.094) | 11.0 |
| | | Birds | 648 | 24.80<br>(p < 0.001) | 1.67<br>(p = 0.196) | 7.1 |
| | | Reptiles | 535 | 29.61<br>(p < 0.001) | 2.10<br>(p = 0.147) | 17.3 |
| | | Amphibians | 179 | 4.99<br>(p = 0.172) | 0.02<br>(p = 0.897) | 3.6 |
| Ecological release<br>Resource limitation<br>Immigration selection | lnRR~ Mainland mass x area + Mainland mass x distance | Mammals | 1038 | 38.36<br>(p < 0.001) | **11.45<br>(p = 0.003)** | 12.0 |
| | | Birds | 648 | 28.66<br>(p < 0.001) | 2.21<br>(p = 0.331) | 8.0 |
| | | Reptiles | 535 | 45.83<br>(p < 0.001) | **12.23<br>(p = 0.002)** | 20.5 |
| | | Amphibians | 179 | 8.93<br>(p = 0.112) | 2.98<br>(p = 0.225) | 5.9 |

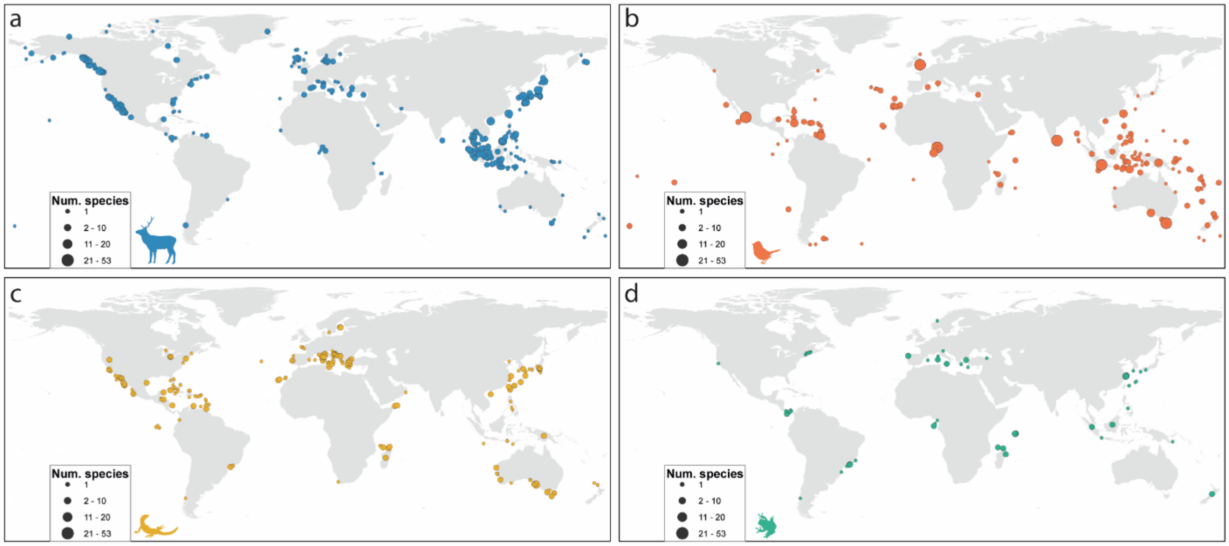


**Supplementary Figure 1.** Location of island populations included in our analyses for (a) mammals (N = 1058, blue), (b) birds (N = 695, orange ), (c) reptiles (N = 547, yellow), and (d) amphibians (N = 179, green). The size of each point indicates the number of species sampled on each island.

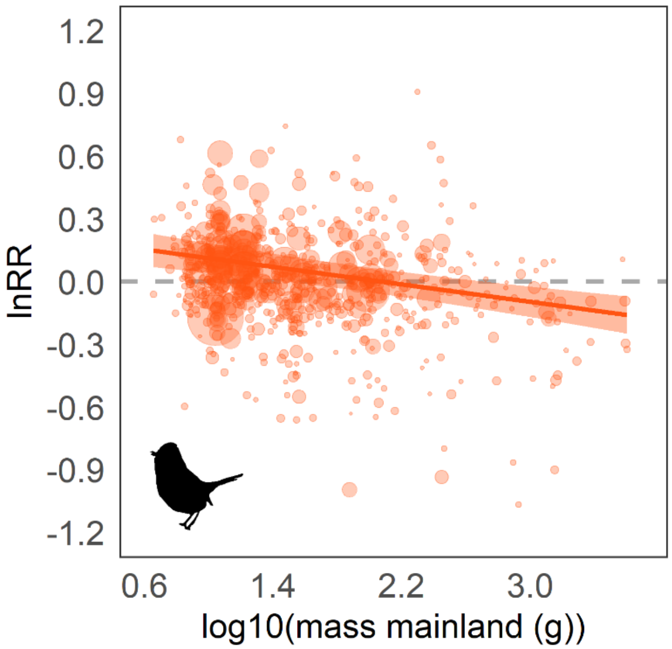


**Supplementary Figure 2. Relationship between *lnRR* (log-ratio between island mass and mainland body mass) and body mass in the mainland for birds when tarsus length was used preferably to convert to body mass equivalents using allometric relationships (Supplementary Table 2). The model was fitted using multi-level, mixed-effects models with mainland body mass as moderator, and observation-level ID, source, species and phylogeny as random effects. *lnRR* > 0 indicates gigantism; *lnRR* < 0 indicates dwarfism; and *lnRR* = 0 indicates stasis (no shift in body size from mainland to island populations). The size of the points represents the weight of each paired island-mainland ratio in the model according to sampling variance. Model estimates are intercept = 0.216 (95% CI: 0.117 – 0.315) and slope = -0.104 (95% CI: -0.145 – - 0.064).**

#### Appendix 1 – Databases searched within WOS

Using our search string, we searched Web of Science Core Collection, including the databases covered by the following subscriptions:

- Science Citation Index Expanded (SCI-EXPANDED) --1900-present
- Social Sciences Citation Index (SSCI) --1956-present
- Arts & Humanities Citation Index (A&HCI) --1975-present
- Conference Proceedings Citation Index- Science (CPCI-S) --1990-present
- Conference Proceedings Citation Index- Social Science & Humanities (CPCI-SSH) --1990-present
- Book Citation Index– Science (BKCI-S) --2005-present
- Book Citation Index– Social Sciences & Humanities (BKCI-SSH) --2005-present
- Emerging Sources Citation Index (ESCI) --2015-present
- Current Chemical Reactions (CCR-EXPANDED) --1985-present
- Index Chemicus (IC) --1993-present

### Appendix 2 – Structured Summary

**Background**
Island faunas can be characterized by gigantism in small animals and dwarfism in large animals, but the extent to which this so-called 'island rule' provides a general explanation for evolutionary trajectories on islands remains contentious.

**Objectives**
We assessed patterns and drivers of body size evolution across a global sample of paired island-mainland populations of terrestrial vertebrates, and evaluated the influence of physiographic, climatic and ecological factors on insular size shifts.

**Data Sources**
We collected data from articles included in a recent assessment of the island rule and previous compilations included in previous studies of reptiles, mammals, and birds, tracing original data sources when possible to extract original measurement data. To avoid the widespread author- or publication-biases detected in previous studies, we sampled body size measurements from published studies that did not assess the island rule per se, or – in the case of birds – also from original morphometric data collected from museum and live specimens. Overall, we included information retrieved from peer-reviewed articles, live and museum specimens, and expedition reports.

**Study eligibility criteria, participants, and interventions**
We included a study if it contained relevant measurement data (i.e. for insular populations or, preferably, for both insular and mainland populations). Unpaired insular populations were matched when possible by performing species-specific searches of adjacent mainland populations in WOS and Google Scholar. Any study reporting morphometric measurements in insular populations was included regardless of whether it tested the island rule or not, thereby avoiding comparisons that were based on species with well-known extreme body sizes only. We also included museum and live bird specimens that were for the first time measured and used to test the validity of the island rule in this study. We discarded studies that reported morphometric data using metric that could not be converted to body mass equivalents using allometric models. We only focused on extant taxa and species that were not recently introduced or invasive. We assembled a global dataset of 2,479 island-mainland comparisons for 1,166 insular and 886 mainland species of terrestrial vertebrates, including mammals (1,058 island-mainland comparisons), birds (695 comparisons) reptiles (547 comparisons) and amphibians (179 comparisons) spread over the globe. In total we included morphometric measurements of 154,875 mainland and 63,561 insular specimens from species covering a wide range of average body masses (0.18–234,335 g). Insular populations in our dataset inhabit a diverse array of islands with different sizes (0.0009–785,753 $km^2$), spatial isolation (0.03–3,835 km from mainland) and different climates.

**Study appraisal and synthesis methods**
Studies were included if they reported measurement data as means, together with sample size and, when possible, a measure of variability (SD, SE, CI). We excluded comparisons that were not supported by taxonomic or phylogenetic evidence, or that were not biogeographically possible. We also excluded cases when only one individual was measured in either mainland or island and thus the SD was zero. We used phylogenetic meta-regression models to assess patterns and drivers of body size evolution across a global sample of paired island-mainland populations of terrestrial vertebrates. We included as random effects Source, Species, Phylogeny and Observation.

**Results**
We show that 'island rule' effects are widespread in mammals, birds and reptiles, but less evident in amphibians, which mostly tend towards gigantism. We also found that the magnitude of insular dwarfism and gigantism is mediated by climate as well as island size and isolation, with more pronounced effects in smaller, more remote islands for mammal and reptiles.

**Limitations**
Our analyses focused solely on extant species for which we could gather data on measurement error and sample size (essential for meta-analyses). The widespread extinction of large species in islands, including dwarf morphotypes of large species such as insular elephants in Sicily and the Aegean islands, may have masked the pattern, making it harder to detect a signal. This suggests that analyses based on present-day patterns may somewhat bias our perception of the rule and scaling coefficients, and that including extinct species would strengthen the signal that we already report for extant species.

**Conclusions and implications of key findings**

We conclude that the island rule is pervasive across vertebrates, but that the implications for body size evolution are nuanced and depend on an array of context-dependent ecological pressures and environmental conditions. Further meta-analyses are needed to assess the consistency of the 'island syndrome' in terrestrial vertebrates.

### Appendix 3. PRISMA Checklist

| Section/topic | # | Checklist item | Reported on page # |
|---|---|---|---|
| **TITLE** | | | |
| Title | 1 | Identify the report as a systematic review, meta-analysis, or both. | NA, not included in the title |
| **ABSTRACT** | | | |
| Structured summary | 2 | Provide a structured summary including, as applicable: background; objectives; data sources; study eligibility criteria, participants, and interventions; study appraisal and synthesis methods; results; limitations; conclusions and implications of key findings; systematic review registration number. | Yes, it is included as part of the Supplementary materials, Appendix 2. |
| **INTRODUCTION** | | | |
| Rationale | 3 | Describe the rationale for the review in the context of what is already known. | 3-4 |
| Objectives | 4 | Provide an explicit statement of questions being addressed with reference to participants, interventions, comparisons, outcomes, and study design (PICOS). | 4, no explicit reference to PICO design. |
| **METHODS** | | | |
| Protocol and registration | 5 | Indicate if a review protocol exists, if and where it can be accessed (e.g., Web address), and, if available, provide registration information including registration number. | NA |
| Eligibility criteria | 6 | Specify study characteristics (e.g., PICOS, length of follow-up) and report characteristics (e.g., years considered, language, publication status) used as criteria for eligibility, giving rationale. | 12-13 |
| Information sources | 7 | Describe all information sources (e.g., databases with dates of coverage, contact with study authors to identify additional studies) in the search and date last searched. | 12-13 |
| Search | 8 | Present full electronic search strategy for at least one database, including any limits used, such that it could be repeated. | 12, Appendix 1 |

| | | | |
|---|---|---|---|
| Study selection | 9 | State the process for selecting studies (i.e., screening, eligibility, included in systematic review, and, if applicable, included in the meta-analysis). | 12-13 |
| Data collection process | 10 | Describe method of data extraction from reports (e.g., piloted forms, independently, in duplicate) and any processes for obtaining and confirming data from investigators. | 14-16 |
| Data items | 11 | List and define all variables for which data were sought (e.g., PICOS, funding sources) and any assumptions and simplifications made. | 12-14, 16-17 |
| Risk of bias in individual studies | 12 | Describe methods used for assessing risk of bias of individual studies (including specification of whether this was done at the study or outcome level), and how this information is to be used in any data synthesis. | 18 |
| Summary measures | 13 | State the principal summary measures (e.g., risk ratio, difference in means). | 15-16 |
| Synthesis of results | 14 | Describe the methods of handling data and combining results of studies, if done, including measures of consistency (e.g., $I^2$) for each meta-analysis. | 17-20 |



| Section/topic | # | Checklist item | Reported on page # |
|---|---|---|---|
| Risk of bias across studies | 15 | Specify any assessment of risk of bias that may affect the cumulative evidence (e.g., publication bias, selective reporting within studies). | 18 |
| Additional analyses | 16 | Describe methods of additional analyses (e.g., sensitivity or subgroup analyses, meta-regression), if done, indicating which were pre-specified. | 18-19 |
| **RESULTS** | | | |
| Study selection | 17 | Give numbers of studies screened, assessed for eligibility, and included in the review, with reasons for exclusions at each stage, ideally with a flow diagram. | 12-13, Supplementary Dataset 3 |
| Study characteristics | 18 | For each study, present characteristics for which data were extracted (e.g., study size, PICOS, follow-up period) and provide the citations. | Page 16, See complete datasets in https://github.com/anabenlop/Island_Rule |
| Risk of bias within studies | 19 | Present data on risk of bias of each study and, if available, any outcome level assessment (see item 12). | We included data in our analyses that was omitted in other studies. |
| Results of individual studies | 20 | For all outcomes considered (benefits or harms), present, for each study: (a) simple summary data for each intervention group (b) effect estimates and confidence intervals, ideally with a forest plot. | See Figure 3. Note that all results are presented as meta-regressions. |
| Synthesis of results | 21 | Present results of each meta-analysis done, including confidence intervals and measures of consistency. | 5-7, Supplementary Tables S3-S8 |
| Risk of bias across | 22 | Present results of any assessment of risk of bias across studies (see Item 15). | 5, Supplementary Tables S3-S6, S8, Fig |

| | | | |
|---|---|---|---|
| studies | | | S2 |
| Additional analysis | 23 | Give results of additional analyses, if done (e.g., sensitivity or subgroup analyses, meta-regression [see Item 16]). | 5-7 |
| **DISCUSSION** | | | |
| Summary of evidence | 24 | Summarize the main findings including the strength of evidence for each main outcome; consider their relevance to key groups (e.g., healthcare providers, users, and policy makers). | 7-10 |
| Limitations | 25 | Discuss limitations at study and outcome level (e.g., risk of bias), and at review-level (e.g., incomplete retrieval of identified research, reporting bias). | 10-11 |
| Conclusions | 26 | Provide a general interpretation of the results in the context of other evidence, and implications for future research. | 11 |
| **FUNDING** | | | |
| Funding | 27 | Describe sources of funding for the systematic review and other support (e.g., supply of data); role of funders for the systematic review. | 55 |

*From:* Moher D, Liberati A, Tetzlaff J, Altman DG, The PRISMA Group (2009). Preferred Reporting Items for Systematic Reviews and Meta-Analyses: The PRISMA Statement. PLoS Med 6(7): e1000097. doi:10.1371/journal.pmed1000097

For more information, visit: **www.prisma-statement.org**. Page 2 of 2
